# Convergent information flows explain recurring firing patterns in cerebral cortex

**DOI:** 10.1101/2025.10.05.680498

**Authors:** Domenico Guarino, Anton Filipchuk, Alain Destexhe

## Abstract

Cortical population events, short-lived patterns of neuronal activity that recur with some consistency, are central to sensorimotor coordination. These reproducible firing patterns are often attributed to attractor dynamics, supported by strong mutual connectivity. However, using multi-modal datasets — including 2-photon imaging, electrophysiology, and electron microscopy — we show that these reproducible patterns do not involve strongly interconnected neurons. Instead, we show that cortical networks exhibit hierarchical modularity, with core neurons acting as high-information-flow nodes positioned at module interfaces. These cores funnel activity but lack structural signatures of pattern completion units expected in an attractor network. Using computational models, we find that distance-dependent connectivity is necessary and sufficient to generate the modularity and transient reproducible events observed in cortex. Our findings suggest that cortical networks are instead pre-configured to support sensorimotor coordination. This work redefines the structural and dynamical basis of cortical activity, highlighting the link between modular structure and function.

## Introduction

All animals produce sensorimotor coordination to survive and thrive. At the top of the nervous system hierarchy, cortical population events of temporally correlated neuronal firing^1^ orchestrate this coordination. Cortical events occur already before birth^2^, promote specific behaviours^3,4^, recur under similar stimulus conditions^5,6^, retain their neuronal composition when reoccurring spontaneously^7,8^, and can also be induced by repetitive coactivation^3,8^. A defining characteristic of cortical events is that they show reproducible firing patterns of their participating cells^7,8^. This reproducibility is often viewed as evidence that these firing patterns are the dynamical signature of fixed point^9,10^, limit cycle^11,12,13^, or continuous^14,15,16,17,18^ attractor networks. Attractor networks are powerful for understanding how neural activity can stabilise into specific reproducible patterns. Neurons within an attractor network tend to excite each other with strong recursive mutual connections, creating positive feedback stabilising the network’s activity pattern^18,19,20^. In limit-cycle attractor models, strong mutual connections ensure periodic behaviour by supporting both excitatory and inhibitory feedback, which drives the cycle forward and regulates its periodicity^21,22^. In continuous attractors, strong mutual connections maintain stability across a continuum of states, resisting perturbations and supporting stable representations of continuous variables like spatial position or orientation^17,23,24^. In the present work, we analysed several datasets to assess whether reproducible cortical population events bear the structural and dynamical hallmarks of attractor networks, or whether an alternative mechanism better accounts for their emergence.

## Results

Cortical population events, also known as ensemble, neural assembly, and cell assembly events, are transiently occurring spiking patterns of groups of neurons^7,25,26^. The study of these events involves primarily two in-vivo techniques: 2-photon calcium imaging, and electrophysiology^27^. 2-photon imaging enables simultaneous high spatial resolution recording of neural activity. Electrophysiological recordings provide high temporal resolution data on the firing of neurons. To study cortical population events, we integrated multiple publicly available datasets: 2-photon data sets from the MICrONS project^28,29,30^ (mouse primary visual cortex, VISp), and equivalent experiments from the Allen Brain Observatory^31^, the Goard lab^32^ (mouse retro-splenial cortex, RSC), the Svoboda lab^33^ (mouse anterolateral secondary motor cortex, ALM), and Neuropixels datasets collected simultaneously from all the mentioned areas and more by the CortexLab^27^.

### Reproducible population events throughout cortex

To identify population events and reproducible spiking patterns, we expanded a previously described method^7^. Briefly, population events were identified as peaks of synchronous firing deviating significantly from a surrogate-based threshold (Fig. 1a-b, example from the MICrONS dataset). Neurons firing within an event were listed in an event vector (Fig. 1a top spike raster, bottom firing rate). Each vector binarily encoded the state – active or inactive – of individual neurons. We then used clustering by correlation, an unsupervised technique designed to group neuronal firing rasters without predefined categories (Fig. 1b). Finally, we selected clusters with significant “pattern reproducibility” – i.e. with a within-cluster vector correlation beyond a surrogate-based threshold. Each cluster of population event vectors represented a pattern, with the actual events being pattern occurrences. A large portion of all recorded events, across both 2-photon (VISp: 0.45±0.21, RSP: 0.64±0.61, ALM: 0.70±0.27) and Neuropixels (means of V1/V2: 0.42±0.21, Parietal: 0.37±0.12, Frontal: 0.48±0.13) datasets, formed clusters with within-cluster correlation values (medians between 0.241 and 0.391, see Fig. 1) exceeding those of surrogate controls. Structured recurrence is thus a generic feature of spontaneous cortical activity.

**Fig. 1:**
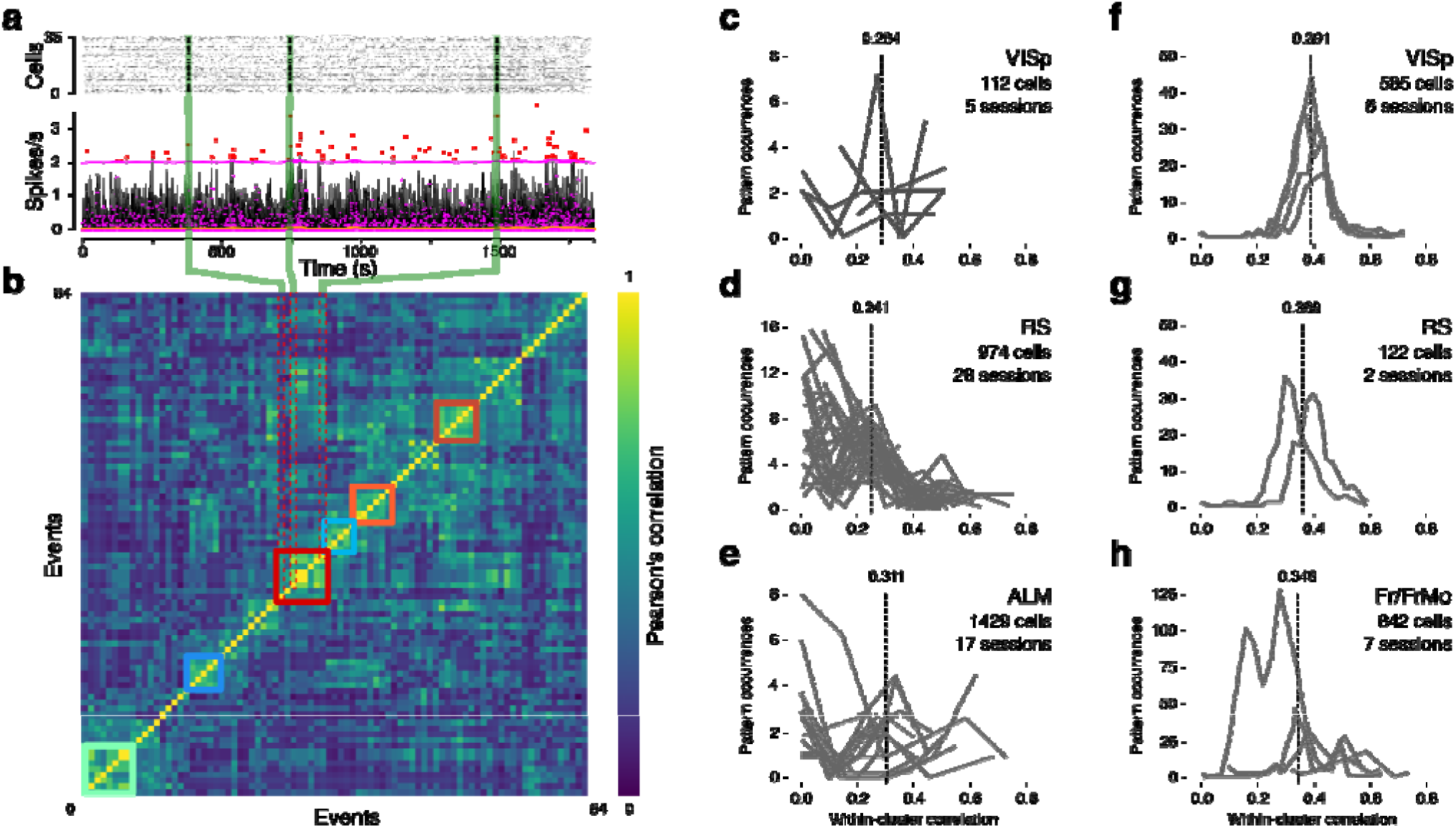
Reproducible activity patterns occur throughout cortex. **a**, (*Top*) Example calcium spikes raster plot from one MICrONS dataset (five available in total). High spike frequency indicate synchronous population events (example green shading). (*Bottom*) Population events were quantified as time intervals where the summed firing rate (black) exceeded a surrogate-based threshold (magenta) based on the baseline (orange). Among all the subthreshold minima (magenta points), those before and after each peak (red squares) are the beginning and end of each event. Each event was assigned a population binary vector (*top*, participating cells as black dots). **b**, Population binary vectors were clustered based on their correlation (colour bar, Pearson’s correlation). Clusters of vectors sharing neurons beyond a surrogate-based threshold were identified (coloured squares, non-significant clusters were omitted). Each identified cluster of population event vectors represents a pattern, with the actual events being pattern occurrences. **c–h**, For each recording session, the number of pattern occurrences vs their within-cluster autocorrelation – reproducibility – was computed. Reproducibility of patterns in the 2-photon (**c**– **e**) and Neuropixels (**f**-**h**) datasets from dorsal (visual primary, VISp), medial (retro-splenial, RS), and frontal cortical (Fr, or antero-lateral motor, ALM) areas for layer 2/3 pyramidal neurons (each panel reports the number of different cells, the number of experimental sessions, and the median reproducibility value, dashed line).

### No pattern completion units in cortex

The above results invite the traditional hypothesis that cortical dynamics, with their reproducible patterns, are akin to attractor networks (and we also confirmed such dynamics, Extended Data Fig. 1). Attractor networks can be tuned to display these dynamics, using biologically plausible Hebbian learning rules^9,19^ to strengthen the connections between co-active units^8,9^. This process can create fixed-point, limit cycle, and continuous attractor dynamics in neural networks^11,18,34,35^. If reproducible cortical activity is the result of attractor dynamics, structural indicators should be found in the cortical architecture. We first tested this assumption in the mouse visual cortex because it has provided connectivity evidence supporting the hypothesis of attractors in the cortex^28,36,37^. For this task, we used the MICrONS dataset^30^. This unique project precisely mapped three million synaptic connections using electron microscopy and co-registered the spiking activity of more than a hundred neurons using 2-photon calcium imaging.

At the cell level (Fig. 2), in line with previously reported statistics^28,36,37^ (Fig. 2a-b), we found few connections between co-tuned cells (Fig. 2c-d). At the population level, our analysis identified 226 significant events. These population events, numerous enough to yield statistical power (88%), exhibited significant specificity for orientation and direction^3,38^ (Fig. 2e), with 27.1% co-tuned cells (having orientation preference within 22.5 degree from the stimulus).

**Fig. 2:**
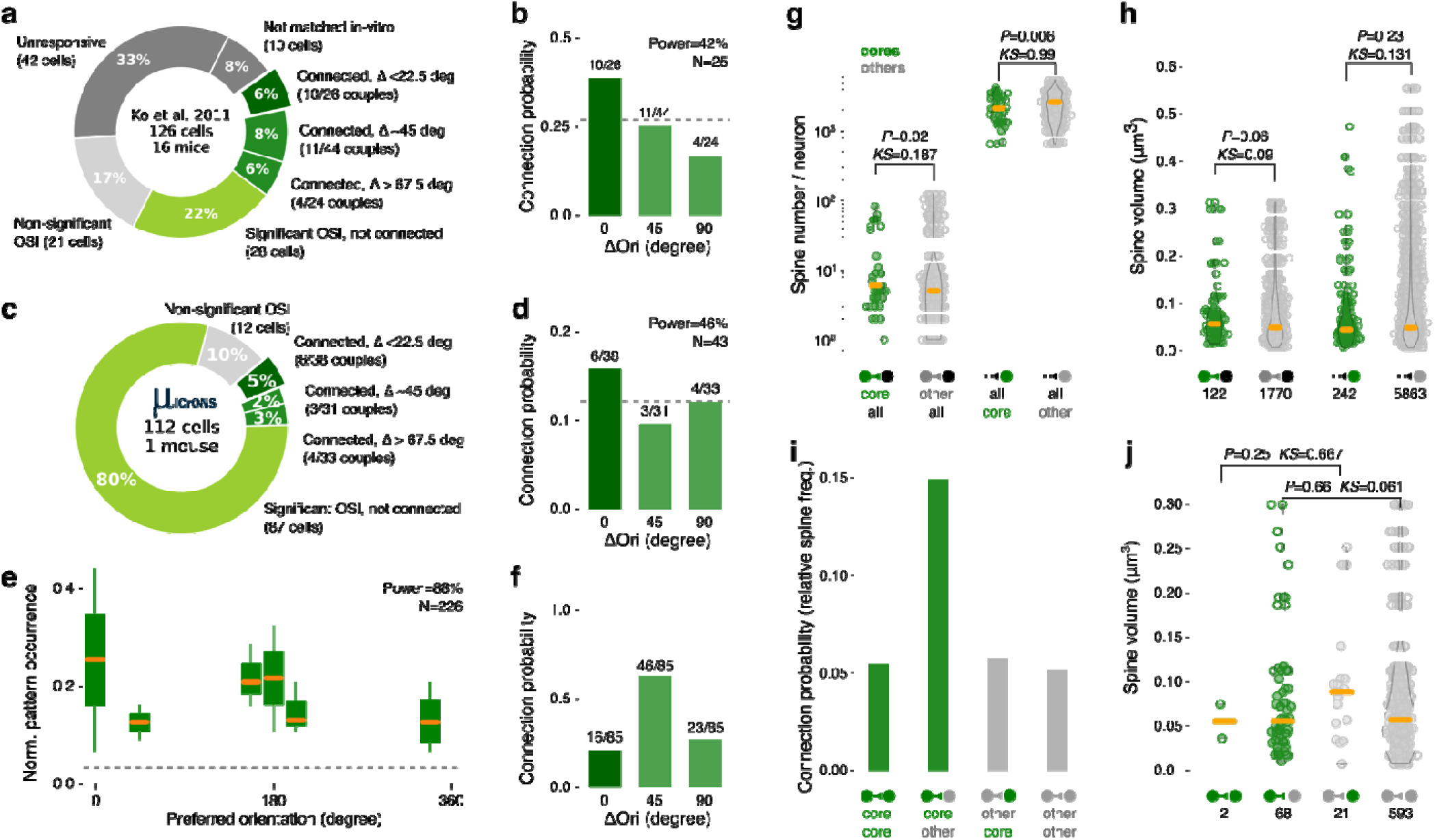
Cell connectivity does not support functional selectivity and pattern reproducibility. **a**, Categorization and number of cells sampled in the original study by Ko et al. 2011. Note the number of cells presenting a significant Orientation Sensitivity Index (OSI), the number of co-tuned and connected cells (green notched slice) against the whole. **b**, Relationship between connection probability and difference in preferred orientation (ΔOri deg <22.5, from 22.5 to 67.5, and >67.5) between cell pairs responsive to grating stimuli (all pairs connection probability=0.27, dotted grey line). The differences in orientation degree values are collected in bins of 45 degrees). The sample for the reported trend is not statistically powerful (42%, N cells). **c**, Categorization and number of cells from the MICrONS dataset. Few co-tuned and connected cells are present. **d**, The relationship between connection probability and difference in preferred orientation for the MICrONS dataset bears no trend nor statistical power (46%, N cells). **e**, In the MICrONS dataset, the reproducible patterns (N=226) were grouped around the horizontal axis (∼0/360 and ∼180 deg, statistical power 88%). **f**, Orientation preference of cells participating in orientation selective reproducible patterns. The majority of participating cells had varying selectivity within 45 degrees from the stimulus. **g**, Number of postsynaptic spines per neuron made by 2-photon imaged cores (green) and other non-core (grey) neurons to all (black) neurons within the EM volume (*left*) and received from neurons within and beyond the volume (*right*). Cores received significantly fewer connections compared to other neurons. **h**, Postsynaptic (proofread) spine volume made by all cores and other neurons to all neurons within the EM volume (*left*) and received from neurons within the volume (*right*). There was no significant difference between spine volumes. **i**, Connection probability for core and other non-core neurons. **j**, Postsynaptic spine volume distributions of cores and others were not significantly different. For **g**, **h**, **j** Kruskal-Wallis significance (P-value) and Kolmogorov-Smirnoff (KS) effect size are reported.

Notably, the cells participating in these events demonstrated diverse orientation preferences, with 72.9% tuned around 45 degrees away from the stimulus, highlighting the ability of population events to recruit cells based on more than individually co-tuned responses (Fig. 2f). Thus, the dataset presented reproducible activities, usually taken as indication of attractor dynamics.

We thus searched for indicators of attractor structure. Reproducible firing patterns do not repeat identically. Only a subset of their member cells – called “core” neurons – reliably take part in many occurrences of the same firing pattern^3,38^. Given this reliable participation in functionally specific events, core neurons are optimal candidates to be pattern completion units^8,38,39^. We singled out core neurons as those participating in most events of significant clusters, and we confirmed core identities using functional correlation40 (Extended Data Fig. 2). The MICrONS 2-photon recordings presented 14 clusters of significant pattern reproducibility (>0.28), with 30 core neurons reliably occurring across events.

Once core neurons were identified, we proceeded with testing the two fundamental attractor-driven assumptions about cortical connectivity: (i) *synapses between cores are more numerous and/or stronger compared to others*, and (ii) *cores have more recurrent connections*^8,37,41^.

To evaluate the first assumption, we used the number and volume of postsynaptic spines^42,43^. When all EM-proofread synapses were taken under consideration, we had evidence (statistical power 92%) to rule out any connectivity preference for core neurons (Fig. 2g and h). Notably for following considerations, cores had a higher probability of having connections to other neurons than to cores (Fig. 2i), only two core neurons presented connections among themselves, and the postsynaptic spine volumes made by core neurons were not bigger than those made by other neurons (Fig. 2j).

To evaluate the second assumption, about the recursivity of circuits comprising cores, we performed several graph theory analyses, considering cells as *nodes* and synapses as *edges* of the network45. To get a global picture of mutual connectivity, we looked at the assortativity *r* – the proportion of mutual edges in a directed graph. We found *r=-*0.08, characteristic of non-mutually connected networks. A detailed analysis of motif distribution46, across all population events of each cluster, including mutual, convergent, and divergent motifs, showed no specific patterns for cores (Extended Data Fig. 3).

Despite lacking direct mutual connections, cores could have been sustained by indirect synaptic feedback, creating cliques of pattern completion units. These possibilities require the existence of more edges, paths, and cycles involving cores than non-cores. We measured the number of cliques, the number and length of shortest paths, and the number of cycles between cores or non-cores. None gave a significant advantage to cores (Extended Data Fig. 4).

To summarise, we found that if we consider core – putative pattern-completion – neurons, their functional and structural connectivity is not biased towards other cores nor co-tuned neurons. These results challenge the applicability of the attractor model to the cortex. But a different approach is possible to explain the presence of population events.

### Cores and the flow of population events

The large number of connections from cores to other neurons (Fig. 2i) suggested their potential centrality in cortical pathways. However, traditional centrality measures (degree, betweenness, and hub scores^45^) showed no significant difference between cores and other neurons (Extended Data Fig. 5). But these static measures ignore dynamics. We thus analysed cortical connectivity as a function of the temporal structure of population events. Events are sequences of cell activations in time (Fig. 3a, *top*) and space (*bottom*). In the MICrONS dataset, we explored whether core neurons influence the flow of activity in the network. The amount of activity that can traverse edges between nodes is measured as “flow”^47,48,49,50^. We measured the minimal flow cut, which is the minimum number of edges that must be removed to disconnect the sub-network defined by each event, going from the first – “source” – firing neuron in the event subnetwork, to the last – “target” – firing neuron. We found that the removal of core connections disrupted the flow between cores as sources and all targets significantly more than other connections (Fig. 3b, p=0.0001, Kolmogorov-Smirnov KS=0.360). This result was confirmed by measuring the PageRank^45^ of the neurons participating in each population event. This algorithm ranks the importance of each node by adding up the importance of every node that links to it, divided by the number of links emanating from it. Thus, PageRank represents a weighted measure of convergence within the neurons in an event. Core neurons had significantly higher PageRank scores than others (Fig. 3c, p=0.00001, KS=0.988).

**Fig. 3:**
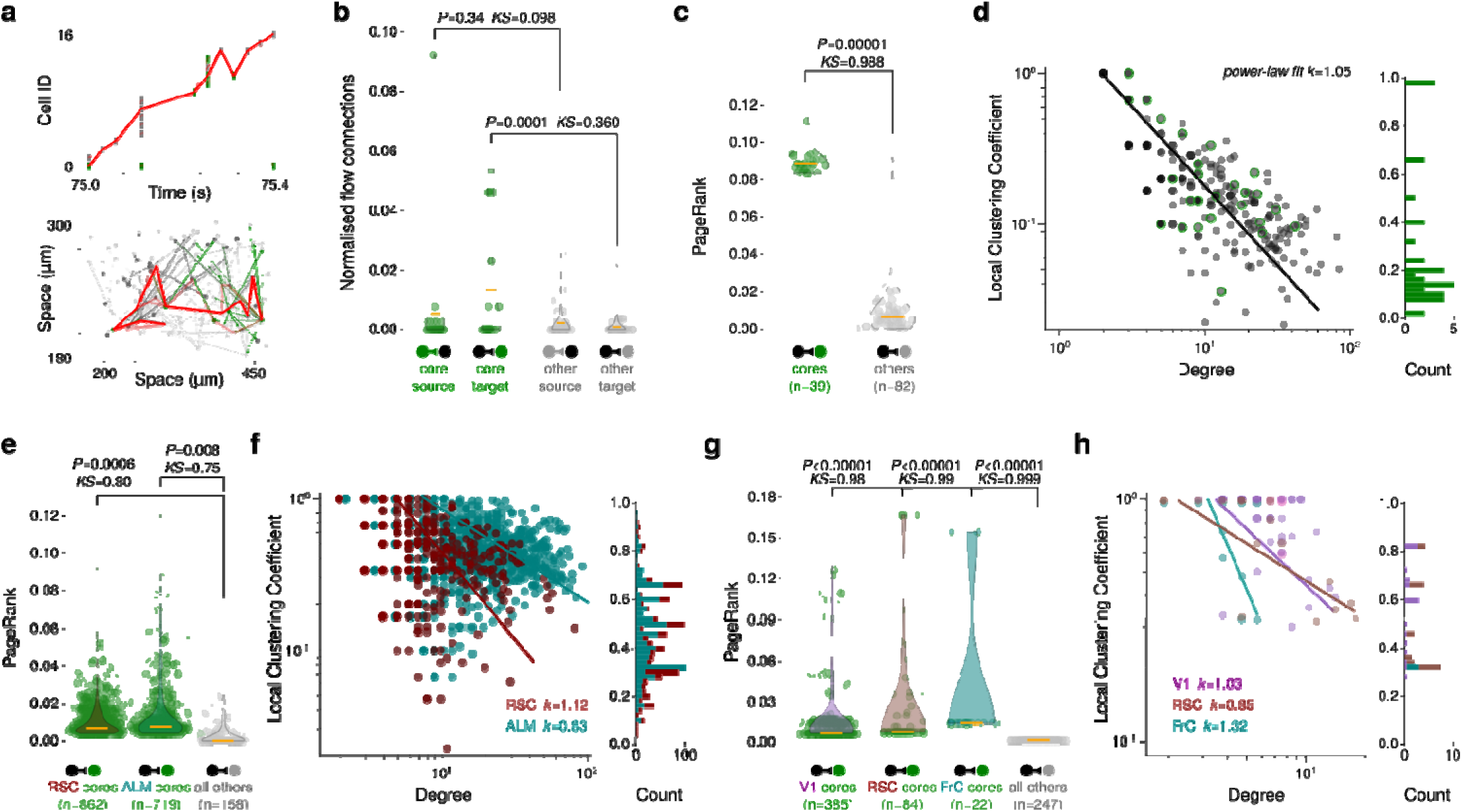
Reproducible patterns run through high information flow nodes of modular networks. **a**, Population events can be studied as flows (red lines, several different flows are possible) in time (*top*), from the first (bottom left) to the last (upper right) firing cell, and in space (*bottom*) over the network. **b**, Neurons can be the source of outgoing and the target of incoming connections. By following all possible pathways between active cells in sequence, we retrieve each cell’s flow capacity between source and target. The removal of connections targeting core neurons (green) disrupted the flow significantly more compared to other neurons (grey). Kruskal-Wallis Significance (P-value) and Kolmogorov-Smirnoff (KS) effect size are reported above the corresponding comparisons (in all following panels). **c**, The PageRank gives a weighted measure of the convergence towards cells participating in events. In the MICrONS dataset, core (green) neurons had significantly higher PageRank scores than others (grey). **d**, High information flow nodes emerge from central positions in modular networks. (*Left*) The log-log linear relationship between the degree – number of connections – and local clustering coefficient – likelihood that neighbours of a specific node are themselves connected – is a measure of network modularity (reported power-law fit). Nodes with high degree and low local clustering coefficients (lower right) are found at the junctions between different modules in the network. (*Right*) The local clustering coefficient distribution of core neurons is skewed towards low clustering values (skewness coefficient 1.53). **e**-**h**, All selected datasets exhibited similar flow, PageRank, and a log-linear relationship, with cores consistently displaying low clustering coefficients (**e**,**f**, 2-photon datasets, **g**,**h**, Neuropixels datasets). Same representations as in previous panels (P-value reported as inequality below 1e-5).

We found that core neurons are not only important for information flow but also critical for maintaining the network’s connectivity. Cores having high PageRank implies that they are typically convergence nodes of the network and central to information flow. Having a high minimal cut indicates that removing these nodes would impact the network’s structural integrity. Together, these measures suggested that convergent connections could funnel incoming activities towards cores. We thus proceeded to better describe the networks embedding core neurons.

### Cores at the gates of hierarchical modules

High information flow nodes typically emerge from controlling positions in modular networks. Nodes within modules have dense interconnections but low flow values. In contrast, inter-module nodes have high flow values due to converging and diverging pathways. A measure of network modularity^40,51^ is the relationship between the number of connections – degree (*d*) – and local connectivity arrangement – clustering coefficient (*C(d)*), the likelihood that neighbours of a specific node are themselves connected. As modules vary in size and interconnectivity, a power-law relationship between clustering coefficient and degree – *C(d) ∼ d^−k^*, with *k∼1* – points to hierarchical modularity. In the MICrONS VISp network, this relationship was *k*=1.05 (Fig. 3d, *left*).

Once VISp modularity was established, we wanted to understand the role of core neurons in it. In the MICrONS dataset, where both activity and structure are known, core nodes with high flow exhibited high degree and low local clustering coefficients (Fig. 3d, *right*), suggesting their role as connectors and gatekeepers between different modules in the network^48,50^.

We confirmed these results outside visual cortex. The other sensory, associative, and motor area datasets analysed exhibited similar flow, PageRank, and log-linear relationship, with cores consistently displaying low clustering coefficients (Fig. 3e, p=0.0006 KS=0.8, and 3h, p=0.00001 KS=0.98). Despite the connectivity being based on functional rather than structural data, we ensured the robustness of connectivity estimation by checking paired short-time (1-5 ms) correlations in the Neuropixels dataset (Extended Data Fig. 6) and confirming that randomly rewiring the MICrONS dataset led to a loss of modularity (Extended Data Fig. 7).

Thus, we found that all inspected cortical networks do not present the structural signature of attractor networks but (i) have characteristics of hierarchical modularity, (ii) have core neurons placed at the interface of modules, (iii) with high information flow and PageRank values. To unify these observations, we propose that distance-dependent connectivity is the fundamental requirement for generating such dynamics.

### Modelling transient reproducible patterns

We offer a mechanistic understanding of how common neural network properties can unify the available evidence to yield transience and reproducibility.

There is an abundance of dissipative models with local connectivity from physics that can inform our understanding of neuronal dynamics in neuroscience^52^. However, these models leave out cortical population events transience and reproducibility, which are usually explained independently using ad-hoc connectivity^53^, dynamically modifying synaptic plasticity^54^, or with additional currents^55^. Based on the strong connectivity assumption of attractor networks, models have been proposed where strong recurrent paths were imposed on randomly connected networks by either additional circuits or increased synaptic strength between arbitrarily selected neurons^18,39,56^. These models yield synchronous firing events corresponding to active sub-population(s). However, the explanatory and predictive power of these models is diminished by their ad-hoc solutions^25^, producing only stereotyped stimulus responses, supporting a narrow set of input□output computations, and requiring strong inputs to exit the attractor state^57,58^.

Here, we present a chain of mechanistic hypotheses: distance-dependent connectivity – decaying in probability with distance – makes hierarchically modular networks, where each module has multiple converging paths funnelling towards high information flow units, which contribute to the reproduction of firing patterns, by limiting the number of possible trajectories. We wove this hypothesis by threading known facts. Distance-dependent connectivity creates modules by allowing predominantly short-range, and few long-range, connections^59,60^. Such modular structure has been shown to occur between the cells of mouse^40^, monkey, and human^61^ cortices. Crucially, in hierarchically modular networks, each module forms a convergent-divergent connectivity motif^51,62,63^, arranged to funnel long and short-range activities toward the input/output nodes, characterised by high flow values^50,64^. In a spiking network, these neurons would become more frequently active leading to reproducible events.

We tested our hypothesis in spiking models by systematically varying the range of connection distances between neurons. To perform the tests, we developed a biologically informed, yet minimal, model of a cortical network of Adaptive Exponential Integrate-and-Fire model neurons65. In building our model, we established connections for each neuron by randomly sampling its neighbourhood with probability exponentially decreasing for increasing distances (Fig. 4a). We chose 9 connectivity ranges evenly spaced between 22 and 90 µm, following the EM distribution of connection distances (Table 2, and Extended Data Fig. 8).

**Fig. 4:**
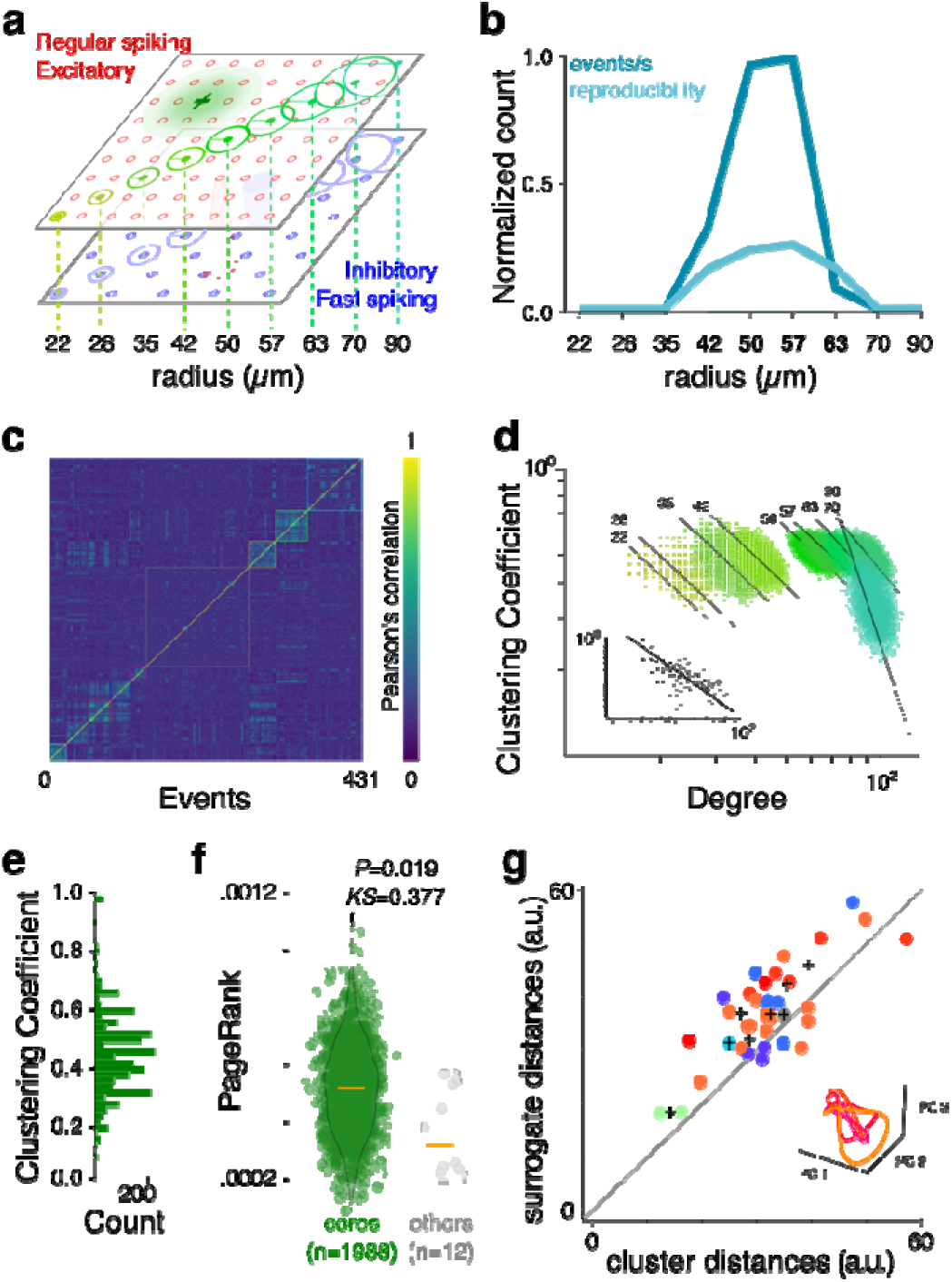
Distance-dependent connectivity creates high-flow core units. **a**, The models were made of excitatory (red) and inhibitory (blue) neurons on a 2D grid (100x100 µm), with a 10 µm step between neurons (10000 neurons in total). Within biological limits, 9 connection range radiuses were tested. **b**, Summary normalised number of events (dark blue) and reproducibility scores (light blue) peaked in the same connection ranges (42-63µm) as the MICrONS data. **c**, Population binary vectors were clustered based on their correlation (colour bar). Clusters of vectors sharing neurons beyond a surrogate-based threshold were identified (coloured squares). **d**, All connection ranges resulted in hierarchically modular networks – i.e. having a power-law relationship between degree and local clustering coefficient of their neurons – (inset, MICrONS data). **e**, The local clustering coefficient distribution of core neurons is not skewed towards low clustering values (skewness coefficient 0.86) but is normally distributed towards low values (mean=0.41). **f**, Core neurons had significantly higher PageRank scores than others (Kruskal-Wallis significance and Kolmogorov-Smirnoff effect size are reported). **g**, The set of simulated population event trajectories (examples in the inset) were more similar to one another than surrogate trajectories (as in Fig. 4d). The multi-dimensional (Hausdorff) vector distance between population event trajectories against that of surrogate.

For all ranges, we tested the presence of population events and cores with flow statistics comparable to experimental results. Over the 9 ranges, only 4 produced synchronous irregular population events. Only to these 4 we applied the same core analysis as above. For example, the model with connectivity range of 50µm (see Fig. 4b), resulted in 431 significant events. Of these events, 273 made 8 clusters of significant reproducibility (>0.2), with 1988 core neurons with population event duration distribution, cluster reproducibility, cluster coefficient and Page Rank as in the MICrONS dataset (Fig. 4c, and below). When exposed to the same stimulus twice, this network produced the same reproducible patterns (Extended Data Fig. 9).

#### Distance-dependent connectivity creates modules

Within the biologically realistic limits just described, we checked that connection distance is sufficient and necessary to produce modular networks (Fig. 4c). We found that all connectivity ranges produced hierarchically modular networks, with exponent *k* close to 1 (*k*=[1.02, 1.3]), as found experimentally^33^ (Fig. 4d, black inset for the MICrONS data *k*=1.05). Distance-dependent connectivity was also necessary for hierarchical modularity. Beyond the physiological limits (>200µm), modularity disappeared from the model, due to the increased reach of connections and the consequent lack of local clustering, and a version of the same network, with randomised connectivity preserving node degree, presented no modularity (Supplementary Information Fig. 2).

#### Distance-dependent connectivity creates high-flow

For the ranges that yielded hierarchically modular networks, with power-law relationships between clustering coefficient and degree close to those found in the MICrONS data, we inspected the organisation of connectivity within each module. As for the experimental data, model cores had a high degree and low local clustering coefficient (Fig. 4e) showing their belonging to the input/output portions of modules. Model cores were significantly more the sources and targets of flow-cutting edges (Supplementary Information Fig. 1) and they had significantly higher PageRank (Fig. 4e), matching the structural-dynamical measurements of real core neurons (Table 3).

The distance-dependent connection strategy was also necessary. We took the above network and randomly rewired all connections, preserving each neuron’s node degree. This randomly rewired network gave rise to population events, but no reproducible pattern was identified, and consequently no cores (Extended Data Fig. 10). The reason is that by increasing the connectivity range, the number of connections and the number of paths increases such that the funnelling disappears^59,60^. Trying to rescue event reproducibility in random networks, for example by increasing input correlations, led to population-wide oscillations far from biologically realistic regimes (Extended Data Fig. 10d).

## Discussion

We looked at transient and reproducible population events happening in local portions of the cortex. We found that these cortical areas are characterized by hierarchical modularity and convergent information flows. Network models indicated that distance-dependent connectivity is necessary and sufficient to produce hierarchical modularity and transient population events with nodes of convergent information flow.

The substantial uniformity of cortical structure and function^67,68,69^, similar findings in other brain regions^70,71^, the genetic encodability of distance-dependent connectivity^72^ and the early establishment of sensorimotor functional maps during embryonic development^73^, allow us to speculate about the generality of these properties and identify a broader implication of our work. All animals need coordinated sensorimotor activity, a function often attributed to postnatal cortical learning^1,74^. However, evidence also indicates that elements of such coordination are present at or even before birth, suggesting the existence of preconfigured circuit motifs. We therefore propose a shift in perspective: population events may not only result from learning but may also act as its developmental substrate. These recurring events could scaffold the emergence of sensorimotor behaviour by providing a structured initial dynamic, later refined through experience-dependent plasticity^71,75,53^, allowing post-action evaluation mechanisms to shape synapses^76^. During life, this repertoire of patterns awaits future experiences. More specifically, learning may not construct the reproducible population events themselves, if these are pre-wired, but instead solidify the transitions between them. This notion is supported by dynamical systems models in which cortical activity evolves through structured sequences of states, with learning shaping the connectivity that governs these trajectorie^54^. If cortex is pre-wired for recurrent co-activation to enable sensorimotor activities, our models suggest that distance-dependent connectivity allows the cortex to provide coordinated facilitation without prior learning. Two objections might be raised about the results presented above. The first is that the available joint dynamical and structural dataset s limited to the visual cortex; other cortical regions might show stronger mutual connectivity between functionally co-active neurons. However, our findings are consistent with previous studies that fail to detect strong mutual connectivity in cortex, and the visual cortex is often taken as an example of an attractor network for stimulus classification^75–77^. The second objection is that mutual connections between functionally correlated neurons have been observed, therefore the point attractor model is valid. However, our analysis of the MICrONS dataset, and Ko et al. 2011 (see Methods), indicates that such connections are statistically too sparse to form the dense reciprocal motifs required by classical attractor models.

Our study faces several limitations. While 2-photon imaging experiments can localize neurons and infer proximity, (i) the Neuropixels recordings lack spatial resolution sufficient to assess inter-somatic distances. Consequently, we cannot exclude the possibility that some correlated units may in fact be anatomically close and mutually connected. This spatial uncertainty limits our ability to generalize the anatomical conclusions from Neuropixels-based functional data. Future multimodal recordings combining fine-scale spatial resolution with high temporal precision will be essential to fully resolve this issue. The (ii) dependence on a single 2-photon/EM mouse dataset potentially results in a reduced and biased view, and (iii) 2-photon temporal resolution and EM sample size could be improved. Two series of facts mitigate these limitations. The substantial uniformity of measures across cortices and techniques supports the current results. In addition, our model, based on a few conservative assumptions, closely mimics experimental features both structurally and dynamically, and the distance-dependent strategy, adaptable to various cortical areas, invites exploration beyond the exponential sampling distribution (Supplementary Information Fig. 2). In another direction, (iv) synaptic connectivity is dynamic, so static EM data may miss transient connections: core neurons might be functionally connected during events, but such connections could be undetectable without time-resolved sampling. This would require an experimentally difficult strategy to get structural samples for the same tissue at largely different time scales. However, at the two timescales we considered, 2-photon and electrophysiology, the cortical networks produced reproducible activity without any sign of strong connections between cores. Relatedly (v), it is possible that attractor dynamics is slower/faster than the resolution of the available recordings. We used 2-photon and Neuropixels – relatively slow and fast – to address this point, finding converging results.

In conclusion, we provided evidence for high information flows and hierarchical modularity in cortex. These results suggest how the brain’s capacity for coordinated activity and learning is pre-configured by simple intrinsic network structures.

## Acknowledgments

This work has been funded by EC Human Brain Project (HBP, grant agreement no. H2020-945539). We express our gratitude to the MICrONS project, Carandini’s, Goard’s, Svoboda’s labs, and the Allen Brain Institute for the efforts they make and their commitment to publicly releasing their datasets, without which this study would have not been possible. We would like to thank Adrienne Fairhall, Yves Frègnac, and Manuel Mameli for stimulating discussions and critically reading the manuscript. The authors received no specific funding for this work.

## Author contributions

Conceptualization: D.G., A.D.

Methodology: D.G., A.F.

Software: D.G.

Validation: D.G., A.F.

Investigation: D.G., A.D.

Visualisation: D.G.

Funding acquisition: A.D.

Project administration: A.D.

Supervision: A.D.

Writing – original draft: D.G.

Writing – review & editing: D.G., A.D.

## Materials and Methods

We performed the same structural and dynamical analyses over data collected in mouse cortex and in biologically constrained conductance-based spiking models.

### MICrONS data source

The MICrONS project (https://www.microns-explorer.org/) made available, through its website, two datasets, corresponding to the first two phases of the project. The first phase, the one we used, has electron microscopy and functional imaging of the same cortical patch of V1, in a 400×400×200 μm volume with the superficial surface of the volume at the border of L1 and L2/3, approximately 100µm below the pia. Here follows a brief description of the procedures^31^.

### 2-Photon imaging

Functional imaging was performed in a transgenic mouse expressing fluorescent GCaMP6f from the Jackson Laboratories (https://jax.org) and the Allen Brain Institute (see next section on the Allen data source). The complete description of mouse transgenic lines, surgical procedures, implants, 2-photon microscope features, stimulation and recording setups, and protocols have already been published^31^. One important feature and the main difference with the Allen Brain Observatory 2-photon dataset is the recording acquisition frame rate, here at ∼14.8 Hz, resulting in timeframes of ∼67 ms (the Allen dataset has double this resolution, ∼30 Hz). The mouse was head-restrained but could walk on a treadmill during imaging.

### Stimuli

30 one-minute of a coloured-noise with periods of coherent motion of oriented noise. Each one-minute trial contained 16 stationary-moving-stationary blocks, with a different direction presented in each block, pseudo randomly ordered. Each of the 16 orientations was presented during 15 frames, followed by 40-41 frames of pink noise. All orientations were randomly repeated 30 times. All scans were made using the same 16 orientations.

### Electron microscopy

After functional imaging, the mouse was perfused with fixing agents, dissected, and 200 µm coronal slices of the V1 area, where the 2-photon sessions were performed, were taken. Serial sections of 40 nm thickness were cut from the slices and prepared for transmission electron microscopy. Custom modifications to the setup and camera allowed fields-of-view as large as (13 μm^2^) at 3.58 nm resolution. Data was acquired in blocks, aligned through multiple steps, and defects manually detected. Somas, synaptic cleft, and spine detection and partnering were manually performed on a (387 clefts) subset for training and testing machine learning libraries. The rest were performed by machine learning libraries. Finally, cells identified in the 2-photon imaging max projection were manually co-registered with EM-identified somas.

### Allen Brain Institute data source

The Allen Institute made available, through its Brain Observatory, a large dataset of neuron responses from several visual cortical areas and layers, using mouse transgenic lines expressing a Cre/Tet-dependent fluorescent calcium indicator (GCaMP6f), which allows the recording of calcium influx associated with neural activity. The complete description of mouse transgenic lines, surgical procedures, behaviour training, implants, 2-photon microscope features, stimulation and recording setups, and protocols were already published (see Allen VisualCoding_Overview whitepaper in the references). Here we briefly provide the relevant information in the context of our work.

### Cell type and experiment selection

The Allen Brain Observatory contains data collected in several cell types of mouse visual cortices (see Allen VisualCoding_Genotyping whitepaper). Cell-type specific expression of GCaMP6f was conferred by the Cre recombinase and tetracycline-controlled transactivator protein (tTA) under the control of the Camk2a promoter with the Ai93 and Ai94 reporters. To maximise the number of recorded neurons across layers, we selected all Cre drivers specific for pyramidal neurons and layers. Considering that each layer of mouse V1 is roughly 100µm thick (Hage et al. 2022), we were able to design cell-type and layer-specific queries which resulted in the selection of the following experiment IDs (as of 2019/09/10). We selected layer 2/3 planes, to have a dataset comparable to the one from the MICrONS project (query: http://observatory.brain-map.org/visualcoding/search/overview?area=VISp&imaging_depth=175,185,195,200,205,225,250,265,275,276,285,300&tld1_name=Slc17a7-IRES2-Cre,Cux2-CreERT2,Emx1-IRES-Cre), selected experiment ids: [502115959, 501574836, 502205092, 502608215, 510514474, 503109347, 510517131, 524691284, 645413759, 660513003, 704298735, 702934964, 712178511, 526504941, 528402271, 540684467, 545446482, 561312435, 596584192, 647155122, 652094901, 652842572, 653122667, 653125130, 657080632, 661328410, 661437140, 663485329, 679702884, 680156911, 683257169, 688678766] (with exp: 501271265 removed due to https://github.com/AllenInstitute/AllenSDK/issues/66). A total of 32 experiments were selected and downloaded as NWB files. Each experiment contained pre-processed calcium indicator fluorescence movies (512x512 pixels, i.e. 400 μm cortical field of view, sampled at 30Hz), corrected for pixel leakage during scanning, ROI filtered, and neuropil subtracted. Each resulting NWB file consisted of an experiment session (∼90 min) at a given location (cortical area and depth).

All original code of this study has been deposited at https://github.com/dguarino/Guarino-Filipchuk-Destexhe and is publicly available as of the date of publication. DOIs are listed in the key resources table. The MICrONS project dataset is available at: https://www.microns-explorer.org/phase1. The Allen Brain Observatory dataset is available at: http://observatory.brain-map.org. The calls to retrieve the Allen data are listed in the Materials and Methods section. An interactive Jupyter notebook is accessible on CodeOcean (https://codeocean.com/capsule/9782876/tree). The code to run all simulations (complete with Dockerfile to recreate the environment) is available at https://github.com/dguarino/Guarino-Filipchuk-Destexhe-simulations. Any additional information required to reanalyse the data reported in this paper is available from the lead contact upon request.

### Other data sources

To widen our analysis to other cortical areas, and ensure the replicability of results, we used two other sources, from Svoboda’s and Goard’s labs. For the antero-lateral motor cortex (ALM), we used the dataset analysed in the article by Li et al. 2015, available for download on the CRCNS website (https://portal.nersc.gov/project/crcns/download/alm-2). For the retro-splenial associative cortex (RSC), we used the dataset analysed in the article by Franco and Goard 2021, available for download on the Dryad data repository (https://datadryad.org/stash/dataset/doi:10.25349/D99W4T). For the Neuropixels recordings, we used the data from Stringer et al. 2019, available for download on the FigShare data repository (http://dx.doi.org/10.25378/janelia.7739750.v4). Given our interest in the firing patterns, we used the available pre-processed unit firing, where the waveforms have been used to identify the units.

### Calcium spike timing extraction

To infer the action potential timings from the calcium indicator fluorescence of the Allen Brain data source, we used the Python library OASIS. This library for spike inference, which is based on an active-set method to infer spikes, also has routines for the automatic estimation of its parameters. The library is available at https://github.com/j-friedrich/OASIS. As parameters for the deconvolve function, we used *penalty*=1, and we used the automatic estimation of fluorescence variation from the data (parameter *g*). The resulting spike timings obtained with these parameters were verified for adherence to the example deconvoluted data provided by the Allen Brain Observatory ipython notebooks.

### Dynamical analysis

For each data source, we applied the same dynamical analysis. The analysis of synchronous irregular population events has already been developed in MATLAB^TM^ and described in Filipchuk et al. 2022. We reimplemented all analysis steps in Python and tested them against the original analysis. Briefly, single-cell activity was described as a cell binary vector, where the number of elements was equal to *T*, the number of time frames during the recording (frame duration was ∼30ms). A value of 1 was assigned at the time frames corresponding to the onset of each spike, and the cell vector was zero otherwise. The number of cell vectors was equal to the total number of recorded neurons in the field of view, *N_e_*, resulting in a *T*x*N_e_* spiketrain matrix where columns corresponded to the time frames and rows to the neurons. Each column was summed to give the number of neurons coactive at every time frame from which the instantaneous population firing rate could be deduced. As is visible from sample raster plots (Fig. 1a), there were periods where spikes were much more synchronised across the population, which could be interpreted as population events, departing from the fluctuations of an asynchronous population spiking process. To identify whether the population events were above chance coincidence, and determine their exact start and end, we applied the following algorithm. (**1**) We created 100 surrogate *T*x*N_e_* spiketrains matrices by reshuffling each neuron’s inter-spike interval. The average 99 percentile (several other percentiles can be tested in the available IPython Jupyter notebooks) across time frames was then calculated and a firing rate threshold for event significance was obtained by adding this average value to a local baseline estimate that aimed at correcting slow fluctuations of background firing rate. The baseline estimate was generated using an asymmetric least square smoothing algorithm from Eilers and Boelens (2005), with parameters *p=0.01* for asymmetry and λ*=10^8^* for smoothness. (**2**) Experimental population firing rate trace was smoothed using Savitzky-Golay filtering, with order=3 and frame window=7, to get rid of non-essential peaks. Local maxima for the smoothed curve that were above the previously defined threshold were retained (Fig. 1b, inset, red square). The adjacent local minima around each of the local maxima were identified (Fig. 1a). Their time frames were taken as the beginnings and ends of population events (Fig. 1a, limits of the green vertical band). (**3**) The time interval between the beginning and end of each population event defined its duration. We described each population event as a binary vector of length equal to the total number of neurons in the recorded field of view (*N_e_*). We initialised all vector elements to *0*, and then we indicated the participation of a cell in the event, spiking at least once over its duration, with a coordinated *1* (Fig. 1a, left). (**4**) Pearson’s correlation matrix for ongoing and evoked events was constructed by computing the Pearson correlation coefficient between all binary vectors. In plotting the matrix, the colour reflected the degree of correlation with a colourmap ranging from 0 to 1. (**5**) Population events sharing a substantial number of neurons, i.e. having a significant degree of correlation, were clustered together (using agglomerative linkage clustering with the farthest distance based on correlation metric, from the Python *scipy.clustering* package), and the correlation matrix could be permuted according to the clustering order. (**6**) The correlation between population events might be simply due to the finite number of recorded cells. We, therefore, defined a significance threshold for intra-cluster correlation, established using surrogate events. We generated *100*x*N_e_*surrogate event vectors (with the same number of cells per event of the originals, but randomly chosen cells, without replacement). We then performed correlation-based clustering (as in point 5) of the resulting surrogate clusters, and we used the 95 percentiles of the correlation coefficient of these clusters as a correlation threshold (other values in the available IPython Jupyter notebooks). For each population *i*, we extracted three figures: □number□of□events, total□number□of□clusters, mean cluster□size (mean number of events per cluster) and computed the 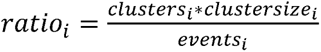 (**7**) To quantify the pattern reproducibility of events within a cluster, i.e. the number of cells shared across events in a cluster, the correlation was calculated across all events of each cluster. (**8**) Core cells were identified as those participating in a range of thresholds (from 55%, barely above no difference in participation, to 95% in the available IPython Jupyter notebooks) of the events within each cluster. Relaxing this requirement to allow for a more participative definition of cores changed the statistics but not the trends presented in Fig. 1. Once identified, we verified that the average firing rate of each core was higher within than that outside its events, discarding those with unspecific firing. In the MICrONS dataset there are 5 2-photon scans, corresponding to different depths. In the first scan, we found 3.86±1.96 (min 0, max 6) cores per cluster, 31.14±1.96 (min 29, max 35) others per cluster. In the second and third scans (grouped to preserve statistical significance), cores per cluster: 1.00±1.41 (min 0, max 3), and others per cluster: 20.00±1.41 (min 18, max 21). In the fourth scan, cores per cluster: 1.50±0.50 (min 1, max 2), others per cluster: 20.50±0.50 (min 20, max 21). In the fifth scan, cores per cluster: 0.75±0.83 (min 0, max 2), others per cluster: 11.25±0.83 (min 10, max 12). In developing our dynamical analysis for the MICrONS data, we used the available notebook example (https://github.com/AllenInstitute/MicronsBinder/blob/master/notebooks/vignette_analysis/function/structure_function_analysis.ipynb). The whole analysis workflow code, is available on GitHub (https://github.com/dguarino/Guarino-Filipchuk-Destexhe), including also an interactive Jupyter Notebook version on CodeOcean (https://codeocean.com/capsule/9782876), which can take a considerable amount of time to download the data from the online resources.

### Attractor analyses

In experimental data, where no system’s characteristic equation is given, we used two levels of analysis. For a network of size *N*, we defined a *state* vector composed of the activity of *N* units. Each *state* comes from a 2-photon frame, deconvolved into continuous dF/F values for the N neurons.

1. At the *population event level*, we study the cell signature of an event. An event *e* is a set of states. At the population level, a vector is formed as the average of all set states. To determine if multiple events form clusters, here we projected each event into a reduced set of embedding dimensions identified using principal component analysis based on the population vectors. We then checked whether the clustered events above occupy different portion of the reduced-dimensional space (and were coloured accordingly). At this level, an attractor is the clustering of events with an average Silhouette score of over 0.7 (for additional details on this method, see the companion Jupyter Notebooks (https://codeocean.com/capsule/9782876).
2. At the *trajectory level*, we studied the approach to and the dynamics of each event (as in Milnor 1985). Here we study if sequences of frames belonging to events from the same cluster have similar trajectories. To quantify the similarity between event trajectories, we adopted the same strategy as Bruno et al. 2017. If noise is the source of discrepancies in the frame trajectories of events from the same cluster, then their distances in the embedding space should be equal to those between artificially recreated trajectories. We measured how closely their trajectories overlapped in the reduced dimensional space of PCA by using the Hausdorff distance. To establish the significance of these distances, we compared them to the distances between surrogate trajectories obtained by shuffling (10000 times) the original frame distances. We established confidence intervals for the Hausdorff distances between actual and surrogate dynamical trajectories using bootstrapping. Coloured dots indicate distances between pairs of frames in the set of events from a same cluster; colour indicates cluster.

### Saddle-node analysis

First, we computed the time *derivatives* of the neuronal activity data. This was achieved using the central difference method. The derivative for each time point *t* was estimated as 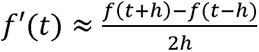, where *h* represents the uniform time step size. For boundary conditions, the first and last derivative values were computed directly, while intermediate derivatives were calculated as the average of adjacent central differences. Then, we computed the Jacobian matrix of the system, which represents the partial derivatives of neuronal activity dynamics. For each time point and pair of neurons *(i,j)*, the partial derivative was computed as: 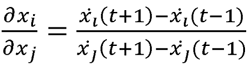, where 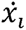 represents the derivative of the activity for neuron *i*, estimated using the Δ*F/F* from the 2-photon data. Finally, we examined the stability of the neuronal dynamics using the computed Jacobians. For each Jacobian matrix, the eigenvalues and eigenvectors were computed using the NumPy ‘eig’ function. The eigenvalues were analysed to identify positive real parts (indicative of instability) and negative real parts (indicative of stability). Time frames exhibiting both positive and negative eigenvalues were classified as saddle node states. These frames were noted for further analysis. For each detected saddle node state, the contributions of individual neurons were assessed based on the eigenvectors associated with positive eigenvalues. Neurons contributing more than 5% to the eigenvector magnitude were identified. Saddle node states were analysed for overlap with predefined event time windows. A permutation test (1,000 iterations) was conducted to assess the expected overlap of saddle node states with events by chance. The proportion of saddle node-contributing neurons overlapping with known cluster cores was calculated for each scan. Observed and expected ratios of saddle node states within events were computed, and statistical significance was assessed using permutation results. Only data from the MICrONS project was used, where the Δ*F/F* data was made available, the other datasets only provided processed spike trains.

### Orientation tuning and cell pairs connection analysis

For the MICrONS dataset, we computed for all 112 2-photon recorded VISp cells, the Orientation Tuning (ORT) as in Ringach et al. 1997: 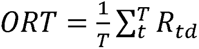 where *T* is the number of trials, *R_td_* is the number of spikes fired during one trial presentation of one stimulus with orientation *d*, and the Orientation Sensitivity Index (OSI) as in Ringach et al. 2004: 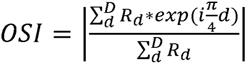, where *R_i_* is the response during stimulus presentation *d*. To test OSI significance, we computed 100 (or 1000) permuted spike trains. We then performed the same connectivity analysis as in Ko et al. 2011. Connection probabilities were calculated as the number of connections detected over the number of potential connections essayed. Having access to all cells simultaneously we sorted all significant orientation-tuned cells by their OSI, then we counted unidirectional/bidirectional connections between all combinations of cells. The connection probability was computed over the total number of couples. Using the computed probabilities, we tested the statistical independence of the connection probability between cells with different orientations (Chi-squared statistical independence). The orientation tuning of population events was computed similarly. See the companion IPython notebook for details on the computations (https://codeocean.com/capsule/9782876).

### Statistical Analysis – general

For all statistical tests, to evaluate the differences between distributions resulting from our measurements of the nominal variables (spine volumes, 1-lag correlations, degrees, betweenness, …) we considered that: the nature of the outcome was continuous, the number of groups was larger than 2 (unless specified, or with unequal measurements), the groups were independent (cores and non-cores groups do not share cells). We tested the distributions’ normality using the D’Agostino and Pearson’s test. For normally distributed variables, we used the One-way ANOVA test. For non-normally distributed variables, we adopted the Kruskal-Wallis test. For some variables, the sample size was large (beyond 100 samples), therefore even small variations could be labelled as significant by the above tests. In these cases, a quantitative measure of the effect size was required. As a measure of effect size, we adopted the Kolmogorov-Smirnov test difference, which is more robust than Cohen’s delta for non-normal distributions (Wilcox and Keselman 2003). For all these tests, we used the Python *scipy* library.

### Power analysis – cell pairs

We evaluated the results from Ko et al. 2011 from the statistical power point of view. From their article: “*The data set contained imaging experiments performed on 16 mice and whole-cell recordings from 126 L2/3 pyramidal cells, 116 of which could be matched to neurons functionally characterized in vivo (see Methods). The rate of connectivity was 0.19 (43 connections out of 222 potential connections assayed), in keeping with previous reports*^6,10^. *Connection probability, synaptic strength and electrophysiological properties of OGB-1-labelled neurons were not significantly different to those recorded in slices from naive age-matched visual cortex that was not injected with OGB-1 AM (connectivity rate 0.18; 25 connected of 143 tested; Supplementary Fig. 2), indicating that dye loading, anaesthesia and prolonged exposure to infrared laser light during imaging in vivo did not alter these parameters. We first examined how connectivity depended on orientation selectivity and on responsiveness to natural movies. Out of the 116 neurons, 77 were responsive to the natural movie, and 79 were orientation selective for grating stimuli (see Methods). Connection probability between orientation-tuned neurons was more than twofold higher than among non-selective and/or non-responsive cells (0.27; 25/94 versus 0.10; 3/31; P=0.050, chi-squared test). The connectivity rate between neurons responsive to the natural movie was significantly higher than among cells non-responsive to the movie (0.28; 30/108 versus 0.04; 2/48; P=0.001, chi-squared test). Taken together, these data indicate that reliably responsive and feature-selective neurons belong to more densely interconnected neocortical subnetworks. We then related connection probability to neuronal preference for the angle and direction of drifting gratings (Fig. 2). For this analysis, we only included pairs in which both neurons were responsive (74/113), orientation selective (orientation selectivity index (OSI) >0.4; 53/74), or direction selective (direction selectivity index (DSI) >0.3; 41/53; see Methods and Supplementary Fig.3a–c).*” Therefore, over 16 mice, 126 cells were recorded in total. 113 cells were recorded both in-vivo and in-vitro. 74 cells were responsive, within which 53 were orientation selective. The actual number of cells found to be co-tuned and connected is not said in the article. But we can compute an estimate, based on the information provided in the Methods and Figure 2a and supplementary Figure 5 of their article. It is said that 26 connection couples were tested for co-tuned cells (and 44 and 24 couples for cells with increasing Δs). Considering that in the Methods is said “*On average we patched 7.9 neurons (range: 2–14) and assayed 13.9 potential connections (range: 2–31) per slice*” (corroborated by Figure 2a and supplementary Figure 5), we can estimate that between 7 and 8 cells were found to be co-tuned and connected (^8^□_2_=28 and ^7^□_2_=21), less than 10 for Δ<67.5□□□, and about 7 cells for Δ>67.5□□□. We wanted to understand if the number of orientation selective, co-tuned, and connected cells found in the study, i.e. 6%, or ∼8 cells of 126, is sufficient to support the claim that local connectivity in the cortex is functionally specific. We took as explicit the usually tacit assumption that cortex in general, and V1 in particular is something like a pattern recognition network, or (fixed-point) attractor network, much like the successful Hopfield model. We aimed to give a simple estimation of the required number of samples to be performed to be able to trust the trends found in the study. The goal was to achieve a desired level of statistical power (e.g., 80%) while maintaining a chosen significance level (e.g., α=0.05). The total number of samples will depend on how many orientation patterns were investigated. Since Ko et al. have 8 distinct orientation patterns, the sampling process need to be repeated for each pattern. We had to take explicit assumptions: ∼5µm thickness for a 2-photon plane of 0.5x0.5 mm, giving roughly 10000 neurons for the network (Kätzel et al. 2011). We assumed a Hopfield-like network of 10000 units with 5% sparse random connections. Given the 5% connectivity, each neuron is connected to roughly 5% of the other 9999 neurons, which is about 500 neurons. We assumed that only a proportion of them have strong connections due to their similar orientation preferences (arbitrarily high 10% = 50 neurons). These percentages need to be applied to each of the 8 orientation patterns. We can provide a simplified estimation of the required sample size for one orientation pattern with the given parameters assuming a standard normal distribution: (i) anticipated mean of connected units (μ0): 50; (ii) standard deviation (σ): 0.025: (iii) significance level (α): 0.05; (iv) desired statistical power (1 -β): 80% (Z = ∼0.845); (v) expected effect size (μ0−μ1): 0.5 (assuming a large effect size results in smaller samples). Our hypotheses, assuming a normal distribution of connections, were: Null Hypothesis (*H0*): *The proportion of significant connections between co-active and co-tuned neurons is equal to or lower than what would be expected by chance* (e.g., 5% due to sparse connectivity). Alternative Hypothesis (H1): The proportion of significant connections is higher than expected by chance. To estimate the required sample size, we used the python power analysis library *statsmodels*. We used the *TTestPower* class to perform a power analysis for the one sample t-test. It calculates the required sample size based on the specified effect size, alpha level, and desired power, assuming we are interested in comparing a sample to the hypothesized population mean (random connections). It resulted in a required sample size for each orientation: 33, total sample size (8 orientations): 267, assuming a standard deviation (sigma) of 0.086. A supplemental post-hoc power analysis was conducted using *G*Power* version 3.1.9.6 (Faul et al., 2007) to determine the actual power tested by the study. Results indicated that, with the sample size of N=25 cells (circa 2 per orientation), at a significance criterion of α=.05, for a one-sample T-test, the power was 42.5% (all this can be replicated in the online IPython Jupyter notebooks).

### Power analysis - population events

We need to determine how many patterns are sufficient to claim that they are functionally linked to the presented oriented patterns. In this case, the effect size is related to the temporal dependencies between the firing patterns and the presented oriented patterns. We counted the number of events per stimulus orientation at increasing lags and normalised it by the total (and visualised the event counts for each orientation, see companion IPython notebook). We used the spread (number of lags with non-zero event counts per orientation, normalised by the total) following a stimulus presentation as a way to quantify the effect size. To know if the occurrence of events meets the required power, we used surrogate data to have a threshold. With a sample size of 226 actual events, a significance of p=0.05, and 10000 random normal surrogate events (a normal distribution assumption, T-test), we obtained a power (spread effect size): 0.8896. Thus, the number of times an ensemble followed an oriented stimulus was sufficient to investigate the (functional) link between events and oriented stimuli. The specificity and significance of pattern occurrences (event occurrence as members of clusters) for oriented stimuli gave 4 significantly orientation-tuned patterns for 7 orientations out of 16 (8 orientation for 2 directions, as in Ko et al. 2011).

### Statistical Analysis – cores and spines

The data released during phase 1 by the MICrONS project concerned one mouse. This limit was our major concern while designing our study. However, our unit of statistical analysis was the number of events. Events are produced by the same cortical tissue overall but by different cells and connections. This implies that we did not set out to measure the same outcome multiple times on the same animal – which would entail an assay variability. Instead, we set out to measure different (quasi-)independent outcomes – which is inherent biological variability (the high level of interconnectivity does not make it completely independent). This is why, in the currently released MICrONS dataset, we consider as statistically legitimate our sample size of n=226 events, i.e. the number of independent observations of our unit of analysis, under a single experimental condition. The goal of our study was hypothesis testing, therefore we designed each analysis as a statistical hypothesis test to produce an exact statistical significance level (*P-value*) with an appropriate effect size. Concerning the attractor features for cores vs non-cores, they are taken to exist if the following hold in the specific direction (one-tailed direction): (1) Core neurons have more numerous synapses between themselves, or towards others, compared to those of non-cores. (2) Spines with presynaptic cores and postsynaptic cores or non-cores are bigger than spines with presynaptic non-cores. **Fig. 2g,h**, We started by assessing whether core neurons made (or received) on average more numerous or bigger spines compared to any other neuron (irrespective of whether them being or not part of the 2-photon imaged dataset). The required sample size, common to both points above (but here calculated for the spine volume), can be computed by considering the following. The study group (“cores”) is compared to the rest of the population (“all”). The primary endpoint is a continuous value (synaptic spine volume, µm^3^). The anticipated mean (µ_0_) of the “others” is 0.05 (based on the recorded average, with σ=0.25). Estimating a higher average for cores of 0.1 (µ_1_), with a type I error probability (false positive rate) of α=0.05, a type II error probability (false negative rate) of β=0.25 (corresponding to a 75% power), the resulting sample size for core spines is *N*=111, since 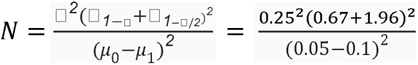. And the smallest count of spines (core-to-all, Fig. 2b, left) is *N*=122. **Fig. 2i,j**, We then turned to the (core-to-core, core-to-other, other-to-core, other-to-other) connectivity of neurons participating in significantly reproducible events, hence limited to both EM and 2-photon imaged neurons. The required sample size, common to both points above, can be computed by considering the following. The study group (“cores”) is compared to the rest of the population (non-cores, or “others”). The primary endpoint is a continuous value (synaptic spine volume, µm^3^). The anticipated mean of the “others” is 0.02±0.12 (based on the recorded average). Estimating a higher average for cores of 0.08, with a type I error probability (false positive rate) of α=0.05, even with a type II error probability (false negative rate) of β=0.3 (corresponding to a modest power of 70%), the resulting sample size for cores is *N*=254. The actual number of 2-photon recorded cells is *N*=112, but the number of cores per event varies between 3 and 8. Therefore, in the status of the MICrONS project, the spine count for cores is underpowered. In order to provide an estimate, we considered four aspects: (a) given that the spine count for each cluster of events with its cores and others is taken independently, many spines will be considered in a repeated measure fashion; (b) the maximum value of the independent variable (spine volume) is >10 times its minimum value (respectively 0.35 and 0.02 µm^3^), allowing a large space of possible values; (c) we lowered the threshold for core identification to a minimum (participation to 60% of the events); (d) we did not differentiate spines based on their point of contact (e.g. axo-somatic, axo-dendritic, axo-axonic, etc).

### Structural analysis

We used the data tables published by the MICrONS project on their website and the referenced publications (see above).

### Postsynaptic spine number and volume analysis

We selected the subset of proofread synapses having as soma root id those identified and co-registered in the 2-photon dataset. After the dynamical analysis was performed (see above), core neuron ids for each event were available and used to query the synapse table, and postsynaptic spine volumes were collected by identified type (core or non-core). The four possible combinations (core-core, core-other, other-core, other-other) were plotted in Fig. 2. Given that different cores participated in different clusters, the number of cores and others differed, and each core (or non-core) neuron could potentially be connected to all other cores (or non-cores) we reported in Fig. 2 the normalised number of spines. Differences in significance and effect size were established using Kruskal-Wallis and Kolmogorov-Smirnov distance, since the distributions were non-normal, as reported below in the statistical methods.

### Synaptic functional efficacy analysis

To estimate functional connectivity, we adopted the same method described by Sadovsky and MacLean (2013). Briefly, an adjacency matrix representing a graph for each experiment was created with nodes being all recorded cells, and directed edges formed according to a single frame lagged correlation (frame duration: 67.4ms) and weighted according to the number of spikes (see Extended Data Fig. 2).

### Graph theory analyses

An adjacency matrix was built by coordinating pre- and post-soma root IDs. All graph analyses were performed using the Python version of the library *iGraph* (Csárdi and Nepusz, 2006). The iGraph library allows motif identification with either 3 or 4 nodes. We used 3-motifs to be able to compare our results with other publications and with the online MICrONS IPython notebooks, and to make computations of the 100 surrogate graphs run in a reasonable time. The comparison of core-based cycles and other-based cycles was performed over an adjacency list and using a function that recursively checked over the list of neighbours starting with each core (or non-core) cell id. In developing our structural analysis for the MICrONS data, we started from the notebook example on the Allen Institute MICrONS GitHub page (https://github.com/AllenInstitute/MicronsBinder/blob/master/notebooks/intro/MostSynapsesInAndOut.ipynb). The whole analysis workflow code is available on CodeOcean as well (https://codeocean.com/capsule/9782876).

### Functional connectivity

To construct the functional adjacency matrix, we analysed binary spike train data for pairwise correlations. The process involved the following steps: (i) cross-correlation, each pair of binary spike trains (*i,j*) was subjected to cross-correlation analysis (with max lag of 1ms in the Neuropixels dataset, 30ms in the 2-photon). Self-connections (*i*=*j*) were excluded by assigning a value of 0 for the diagonal elements of the matrix. The results of the pairwise cross-correlation analysis were stored in a 2D array, forming the functional adjacency matrix. To ensure the sparsity of the adjacency matrix, weak correlations (<0.4) were set to 0. This threshold was based on the approach described by Sadovsky and MacLean (2013).

### InfoMap analysis

To identify structural modules in the MICrONS data and the models, which are directed networks, we used a random walk approach described by Rosvall and Bergstrom (2008). We also used the same algorithm to identify, within each module, the largest connected component and the in-/out-components to the central one.

### Network models

All simulated networks were composed of n=12500 (excitatory/inhibitory neuron ratio of 4:1, i.e. 10000 excitatory and 2500 inhibitory) conductance-based adaptive exponential integrate-and-fire (AdExIF) point neurons. The set of equations describing the membrane potential (1) and spike adaptation current (2) of a neuron are

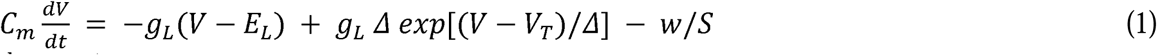

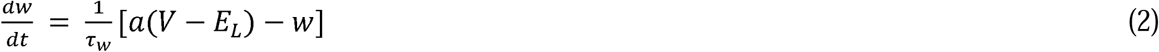

with *C_m_* being the membrane capacitance (nF), *g_L_*the resting (leak) conductance (µS), *E_L_* the resting (leak) potential (mV, which is also equal to the reset value after spike), Δ the steepness of the exponential approach to the threshold (mV), *V_T_*the spike threshold, and *S* membrane area (μm^2^). When *V* reaches the threshold, a spike is emitted, and *V* is instantaneously reset and clamped to the reset value during a refractory period (ms). *w* is the adaptation variable, with time constant τ*_w_* (ms), and the dynamics of adaptation is given by parameter *a* (in μS). At each spike, *w* is incremented by a value *b* (in nA), which regulates the strength of adaptation. An experimentally informed parameter set was chosen (from Zerlaut et al. 2016) and kept fixed, see Table 1 below.

**Table 1.** Parameters of the AdEx Integrate-and-Fire model.

| Parameter | Value | Parameter | Value |
| --- | --- | --- | --- |
| **Regular Spiking (excitatory)** |  | **Fast Spiking (inhibitory)** |  |
| Threshold ($V_T$) | -50.0 mV | Threshold ($V_T$) | -50.0 mV |
| Resting potential ($E_L$) | -65.0 mV | Resting potential ($E_L$) | -65.0 mV |
| Afterpotential reset | -65.0 mV | Afterpotential reset | -65.0 mV |
| Refractory time constant | 5 ms | Refractory time constant | 5 ms |
| Membrane time constant ($\tau_m$) | 15.0 ms | Membrane time constant ($\tau_m$) | 15.0 ms |
| Membrane capacitance ($C_m$) | 0.15 nF | Membrane capacitance ($C_m$) | 0.15 nF |
| Subthreshold adaptation ($a$) | 4.0 nS | Subthreshold adaptation ($a$) | 0.0 nS |
| Spike-triggered adaptation ($b$) | 0.02 nA | Spike-triggered adaptation ($b$) | 0.0 nA |
| Slope factor ($\Delta_T$) | 2.0 mV | Slope factor ($\Delta_T$) | 0.5 mV |
| Adaptation time constant ($\tau_w$) | 500 ms | Adaptation time constant ($\tau_w$) | 500 ms |

### Synapse model

The synaptic connections between neurons were modelled as transient conductance changes. The synaptic time course was modelled as an alpha function with a fast rise followed by an exponential decay:

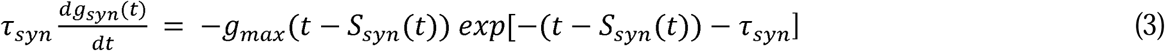

where *syn*∈*{exc, inh}*, *S_syn_(t)=*δ*(t − tk)* are the k incoming synaptic spike trains where i∈{1, .., N} refers to presynaptic neurons and k to the different spike times of these neurons. The synaptic time constants were chosen to be τ_exc_=3 ms and τ_inh_=7 ms for excitation and inhibition respectively. The reversal potentials were E_exc_=0 mV and E_inh_=−80 mV. The synaptic weights g_syn_(t) were set to g_exc_=1 nS for the excitatory conductance, and a balance of g_inh_=5 nS unless stated otherwise. We used a constant delay of 0.5 ms, equal for all connections.

### Network architecture

We simulated a microscope focal plane area of 100x100 µm as a 2D-layer-like network (with periodic boundary conditions to avoid size effects). Neurons were arranged on a grid meant to represent ∼10 µm separation between neurons. Every neuron was sparsely connected with the rest of the network with a connection probability drawn from distance-dependent probability distribution. Each neuron *i* was connected with another neuron *j* with a probability *p_ij_*, depending on their distance *r_ij_*, according to an exponential profile:

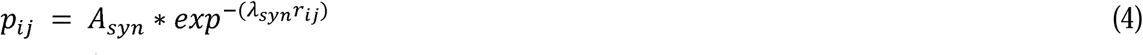

where *A_syn_* is the amplitude of the exponential, varying the number of neurons sampled in the chosen radius *r_ij_*. The amplitudes *A_syn_* were set to have the number of connected neurons roughly match the distributions described by Kätzel et al. 2011, Markram et al. 2015, and Seeman et al. 2018, considering the topological distribution and the excitatory/inhibitory neuron ratio (*A_exc-exc_*=14, *A_exc-inh_*=24, *A_inh-exc_*=14, *A_inh-inh_*=24), resulting in a degree of ∼75 connections per neuron. The decay controller □*_syn_* was set to □*_exc_*=1.2 and □*_inh_*=1.5. The seed to the random number generator responsible for the sampling of the connections was systematically varied, producing N=20 different networks, but following the same type of exponentially decaying distributions. In the tests for the relationships between distance-dependence and flow measures, we systematically varied the amplitudes, lambdas (and correspondingly synaptic weights to keep the conductance balance constant across network configurations). The arrangement of connections was tested for changes in synaptic weights. See below a summary table of modularity measures and population events in the model (Table 2).

**Table 2.** Summary of various measures in models and data.

| connectivity range ($\mu\text{m}$) | power-law fit ($k$) | $R^2$ | normalised bow-tie / tot communities | units / community | population events / s | pattern reproducibility |
| --- | --- | --- | --- | --- | --- | --- |
| 22 | 1.02 | -3.45 | $0.56 \pm 0.2$ | $42.6 \pm 0.6$ | – | – |
| 28 | 1.02 | -0.65 | $0.61 \pm 0.2$ | $49.2 \pm 0.5$ | – | – |
| 35 | 1.03 | 0.15 | $0.68 \pm 0.3$ | $68.3 \pm 0.3$ | – | – |
| 42 | 1.03 | 0.10 | $0.70 \pm 0.1$ | $82.6 \pm 0.4$ | 1.00 | 0.14 |
| 50 | 1.03 | 0.01 | $0.82 \pm 0.3$ | $113.1 \pm 0.2$ | 2.92 | 0.25 |
| 57 | 1.03 | 0.06 | $0.71 \pm 0.4$ | $128.0 \pm 0.3$ | 2.99 | 0.27 |
| 63 | 1.20 | 0.04 | $0.65 \pm 0.2$ | $153.3 \pm 0.1$ | 0.24 | 0.12 |
| 70 | 1.30 | 0.03 | $0.62 \pm 0.5$ | $177.0 \pm 0.4$ | – | – |
| 90 | 3.34 | -1.57 | $0.60 \pm 0.2$ | $253.7 \pm 0.5$ | – | – |
| data | 1.05 | -10.5 | $0.17 \pm 0.1$ | $22.5 \pm 0.6$ | 1.21 | 0.18 |

### Drive

An external population of Poisson drivers was connected to both the RS and FS populations, with uniform random probability drawn from the half-open interval [0,0.01), with a fixed synaptic weight of 0.5 nS.

### Simulator

All simulations were performed using the NEST simulator (Diesmann and Gewaltig 2001) controlled through the PyNN interface (Davison et al. 2009).

### Comparison of model and MICrONS properties

Simulating a network whose only structure arose from distance-dependent connectivity yielded properties comparable to the MICrONS dataset, listed below (Table 3).

**Table 3.** Comparison of model and MICrONS properties.

| MICrONS property | Network model compatibility |
| --- | --- |
| Distribution of distances between connected neurons | An exponential distribution with a single $\square_{\text{syn}}$ value from eq. 4 gives a limited adherence to real distributions. In the Supplementary Information Fig. 2, we tested a normal distribution of $\square_{\text{syn}}$ values and found a better distribution of distance (see also Kätzel et al. 2011, Hage et al. 2022, Seeman et al. 2009). |
| Distribution of neuron degree | The simulator is set to keep the number of connections per neuron constant, this is advantageous to control the conductance-balance, but real neurons present a distribution of degrees. In the Supplementary Information Fig. 2, we also tested the possibility of a degree distributions (see also Turner et al. 2022). |
| Motif distribution | The MICrONS data presents a well-known distribution of motifs (Sung et al. 2005, Turner et al. 2022). The network model reproduces partially motifs distribution (see Supplementary Information Fig. 2). |
| Relationship between Calcium (data) and Sodium (model) spikes | see Supplementary Information Fig. 3 |
| Stimulus specificity of reproducible population events | see Extended Data Fig. 9 |
| The network is hierarchically modular | see Fig. 4d |
| Model core neurons are the target and source of cutting edges | see Supplementary Information Fig. 1 |
| Core neurons are input/output nodes of modules | see Fig. 4e |

## Data Availability

The datasets analysed during the current study are available in the Code Ocean capsule hosted by Nature (https://doi.org/10.24433/CO.9782876.v3)

## Code Availability

The custom code used to pre-process data, conduct analyses, and reproduce all figures is available in the Code Ocean capsule (https://doi.org/10.24433/CO.9782876.v3)

