## Supplementary Information for "Convergent information flows explain recurring firing patterns in cerebral cortex"

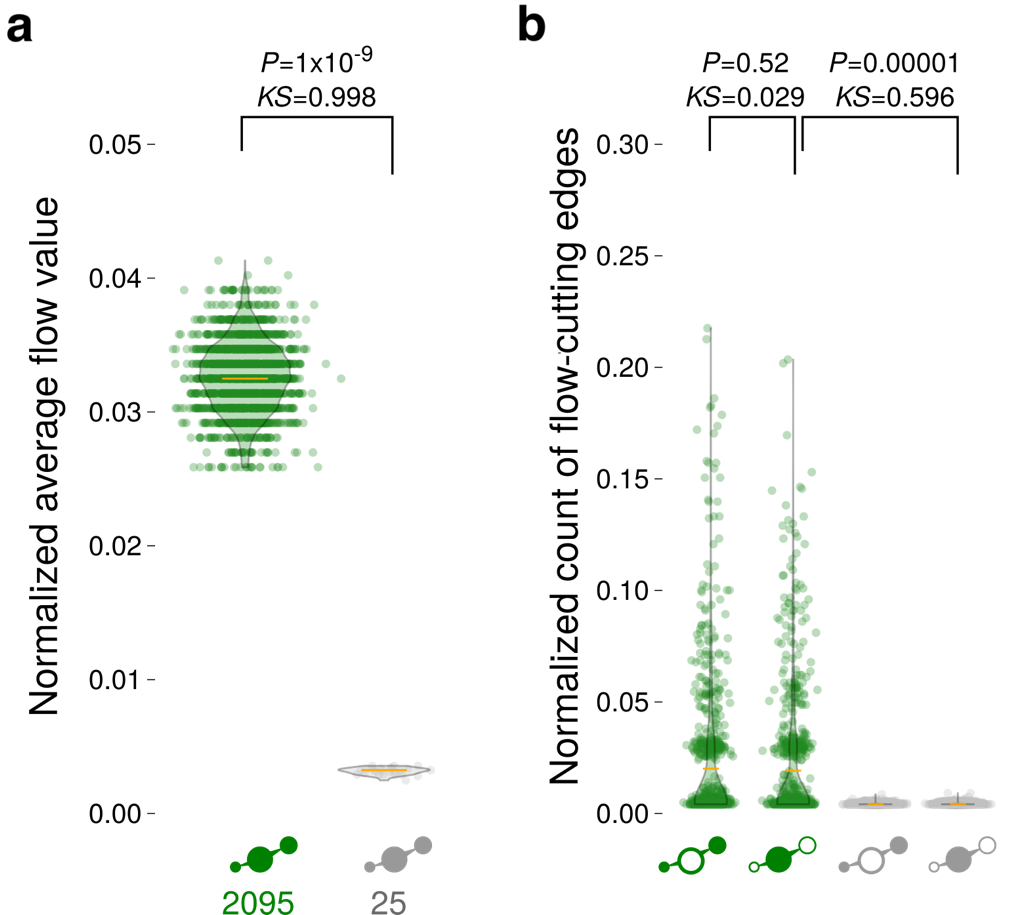

**Fig. 1. Model cores were significantly more the sources and targets of cutting edges. a**, Core neurons sustained significantly more than others the flow of event sequences they participated in (example from range 50µm). **b**, Removing core connections from the network graph stopped more event sequence flows than others.

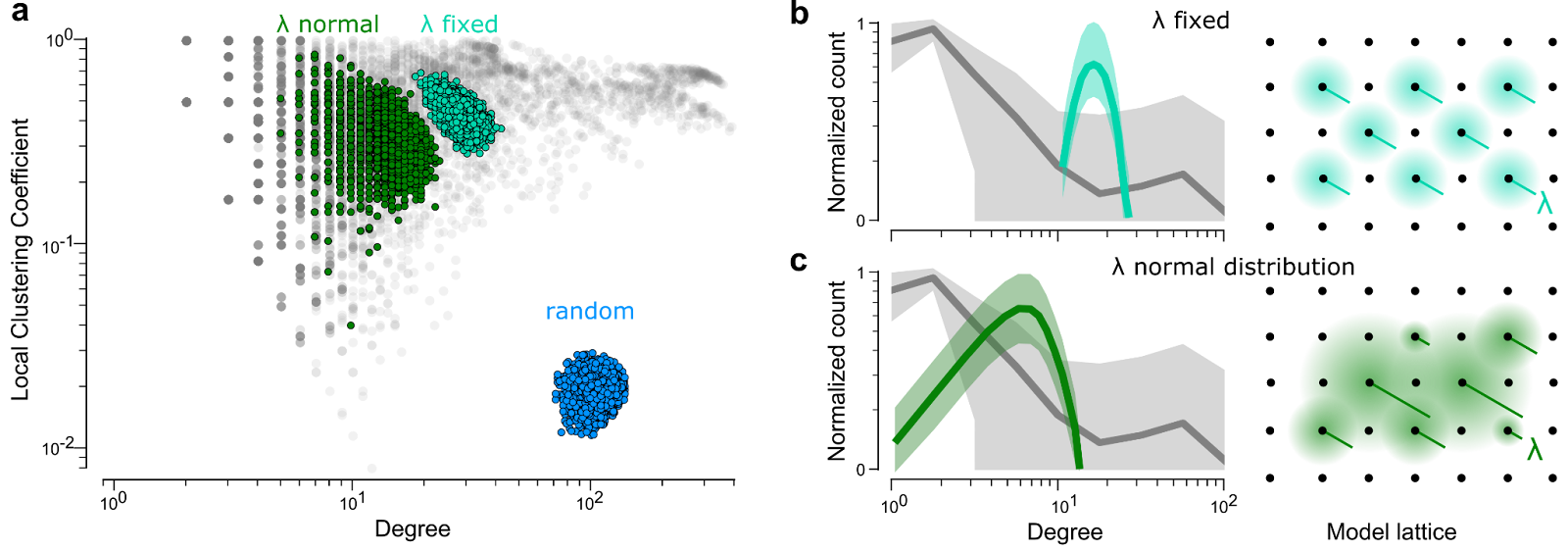

**Fig. 2. Hierarchical modularity of different model connectivity strategies. a**. Different connectivity strategies lead to different relationships between node degree and local clustering coefficient (LCC). Natural networks present a variety of relationships (grey). Completely random connectivity, ignoring the spatial position of units, led to networks in which degree and LCC bear no measurable log-linear relationship and cannot be qualified as modular (blue). Networks in which we used a spatial exponential decay of connection probability between units *i* and *j*, with decaying factor λ (*p_ij_=A_syn_*exp^-(λr)^*) presented log-linear relationships between degree and LCC (see main text and Methods). **b**. Fixed λ for all connections, which is the one used in the models described in the paper, led to compact hierarchically modular networks (cyan, one example) compared to the distribution from natural networks (grey, SEM shaded area). **c**. Normal distributions of λ, which are not described in the paper, led to larger distributions of degree-LCC relationships, stll visibly different from experimental data (green, one example for a normal distribution of λ with mean=1.2 and spread=0.7; grey, same distribution for natural networks).

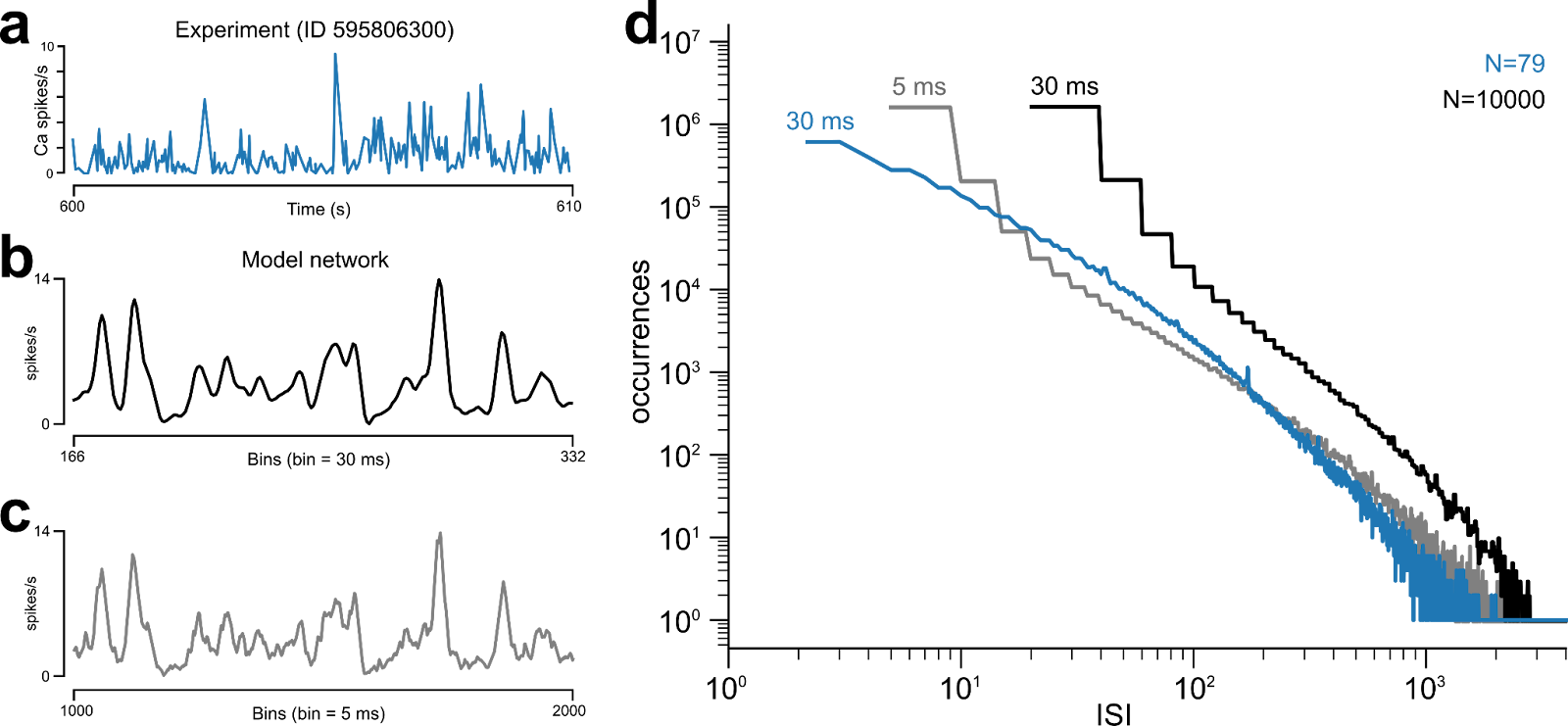

**Fig. 3. Relationship between Calcium (2-photon) and Sodium (model) spikes**. To ensure a good match of modelling population events with the Calcium data, we used the inter-spike interval (ISI) in the firing rates of each recorded neuron (from the much larger dataset of the Allen Brain Observatory, see Methods). We took the experimentally measured rates (**a**, in blue, sampling is ~33Hz), and differently binned spiking activities from the model (**b**, black: 30ms bin, **c**, grey: 5ms bin, all traces are examples). **d**, We then we checked the distribution of experimental and model ISI firing rates. We found that 5ms binning for the model firing best approximated experimental ISIs.
