## Extended Data for "Convergent information flows explain recurring firing patterns in cerebral cortex"

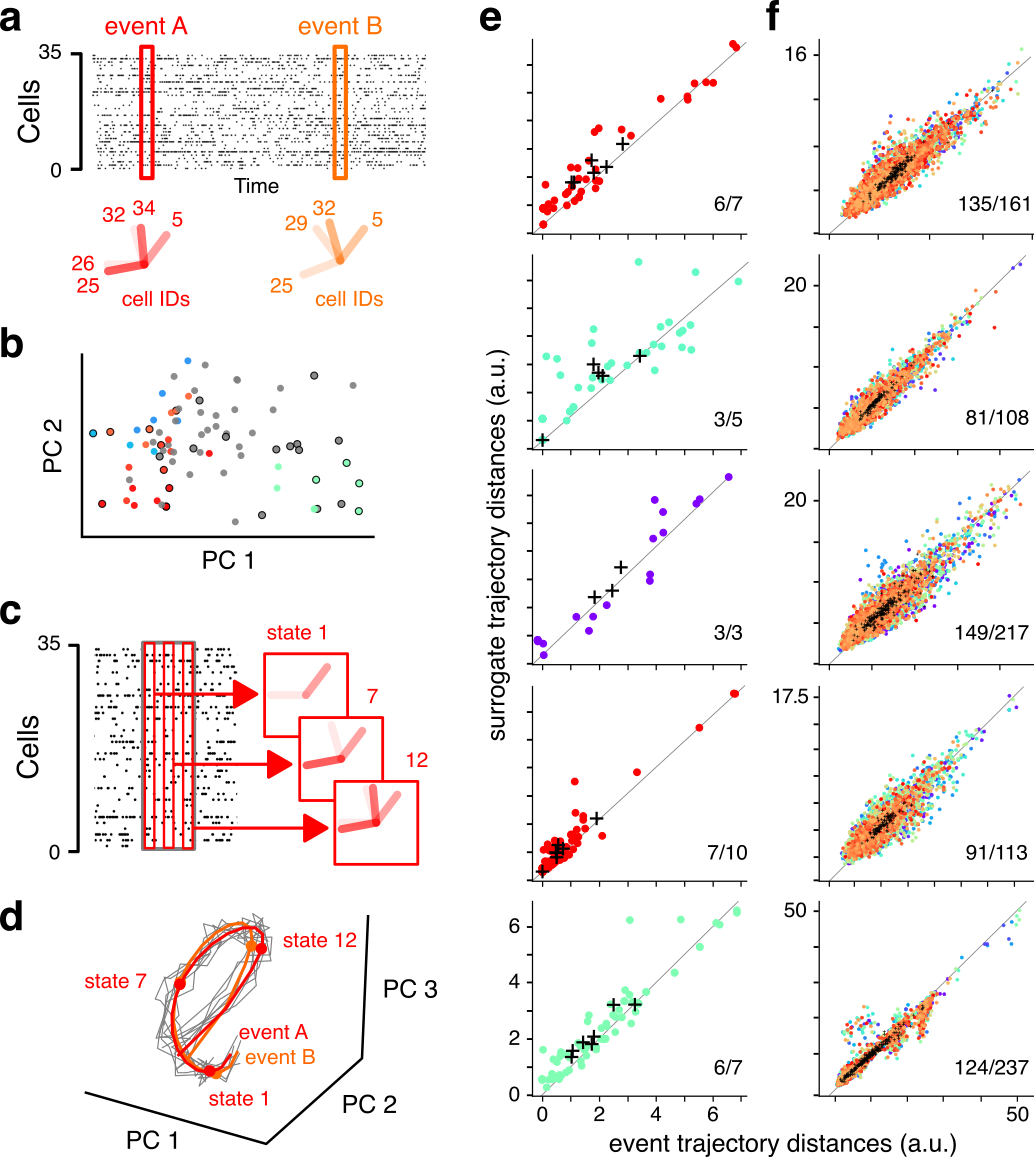

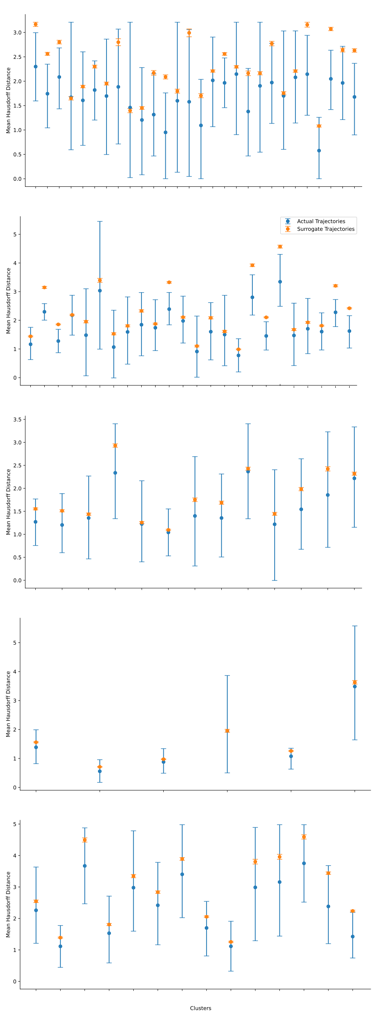

**Fig. 1. Reproducible patterns are transient attractors**. **a**, (*top*) Population events were identified (as described in Fig. 1, two examples A and B). (*Bottom*) The dF/F value of each cell participating in the event can be represented as a radial multidimensional vector defining the state of the system. **b**, An eigenvalue decomposition allows the choice of a suitable reduced set of axes to represent the multidimensional vectors. By looking at their (reduced) state space, the events are grouped in space and coloured as the clusters in Fig. 1b (those occurring during stimulus presentations have black edges). Their Silhouette distance confirmed the clustering (see text). **c**, We can zoom into each event dividing it into smaller states (*left*). Each state is characterised by cell dF/F vectors as in **a** (*right*). **d**, The set of states of an event form a trajectory in the (reduced) state space (red and orange lines). To establish a significant distance, we created surrogate trajectories (grey lines) by shuffling cell vectors, and we measured the multi-dimensional (Hausdorff) vector distance between population event trajectories against that of surrogate trajectories. **e**-**f**, Each event is represented by a dot, coloured by its cluster. Its coordinates are the Hausdorff distances within the cluster and within surrogates (black cross at mean trajectory distance, mean surrogate distance). The majority of cortical recordings presented shorter distances (points coloured by cluster, with means as crosses) between event trajectories compared to surrogates (**e**, events from the MICrONS dataset, **f** events from the Neuropixels dataset, with some confidence intervals shown on the right column the associated notebook).

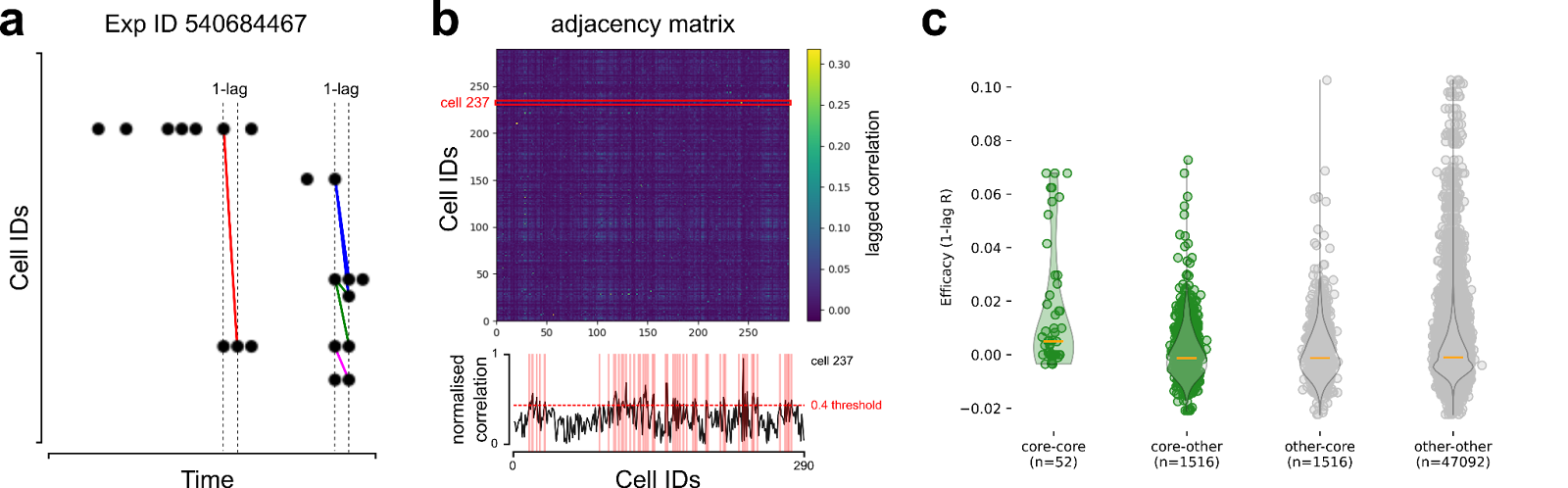

**Fig. 2. Core functional correlation**. **a**, Detail from the Allen Brain Observatory experiment ID 540684467 raster plot showing calcium spikes (black circles). Each row contains the calcium spikes for a cell. The minimal interval between two spikes is the 2-photon imaging time resolution (30 Hz for the Allen Brain dataset). This interval represents the 1-lag correlation interval to identify cells potentially connected. In the figure, only some cells with 1-lag correlated firing were joined by coloured segments to improve legibility. **b**, The vector of 1-lag correlations for each cell is collected into a matrix (top). Weak correlations (threshold=0.4) were not considered for further analysis (bottom, correlations for cell id 237). **c**, Core neurons are those sharing (non-significant, t-score=1.53, *p*=0.12) more high functional correlation across events compared to others.

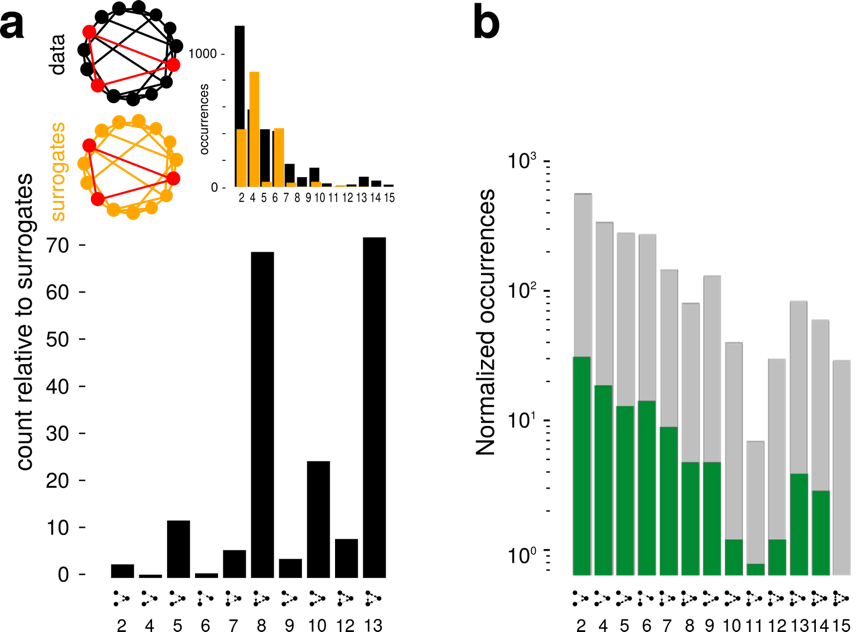

**Fig. 3. No specific motifs for core neurons.** We looked at the distribution of motifs – statistically significant connectivity patterns considering groups of three cells. To understand the significance of motif occurrences, we used the ratio of real vs 100 surrogate degree-matched networks. **a**, Connectivity motifs distribution for all 2-photon recorded neurons (*top*: occurrences count). The ratio followed a known distribution showing an abundance of three-cell mutual motifs. **b**, Normalized motif occurrence distribution for cores and other neurons. Both made a mixture of mutual and non-mutual connections.

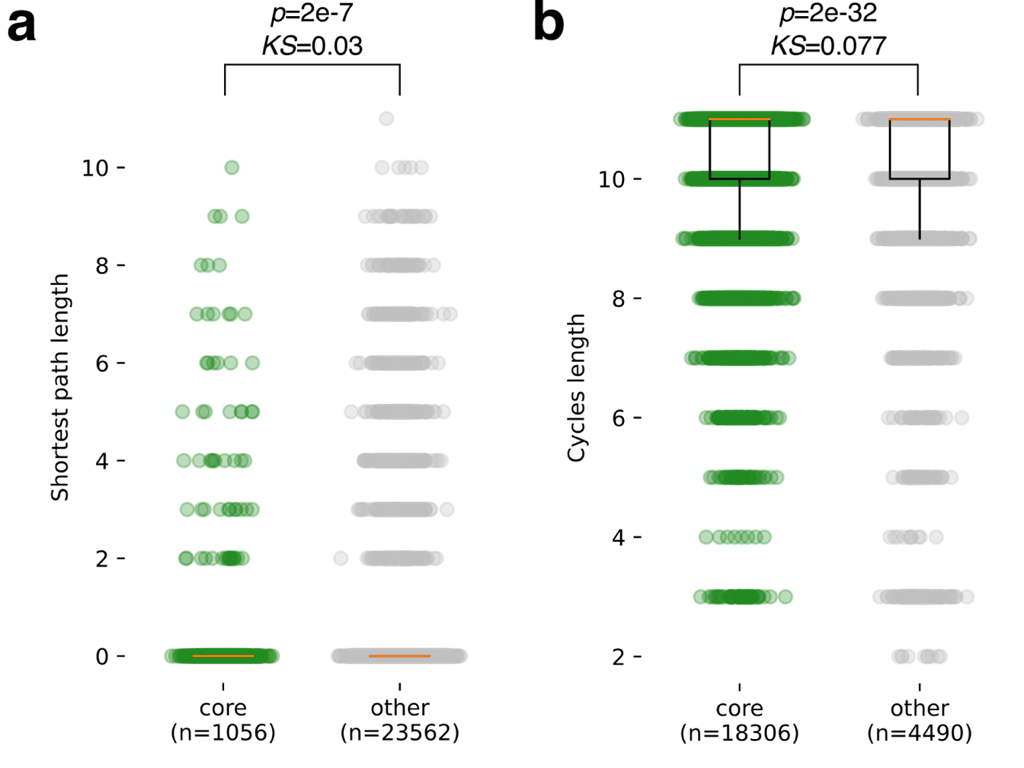

**Fig. 4. Cores are not recursively connected via multi-synaptic feedback**. Even if cores were not directly connected, they could be highly connected via secondary paths, abundant enough to ensure that they are pattern completion units in traditional attractor network terms. **a**, Core-to-core shortest paths were neither more nor shorter than other-to-other paths (0.27±1.19 vs 0.16±0.92 length, Kruskal-Wallis t=26.4 p=2.7e-07, *KS*=0.03). **b**, Core neurons could be part of looped paths, circling back to them, and providing a weak form of recursion. Core-based cycles were more (green, 18306) compared to other-based cycles (grey, 4490). But core and non-core cycles had on average the same length (10.34±1.07 and 9.93±1.76 connections, with minimal effect size KS=0.077), same as the network diameter (*d*=10).

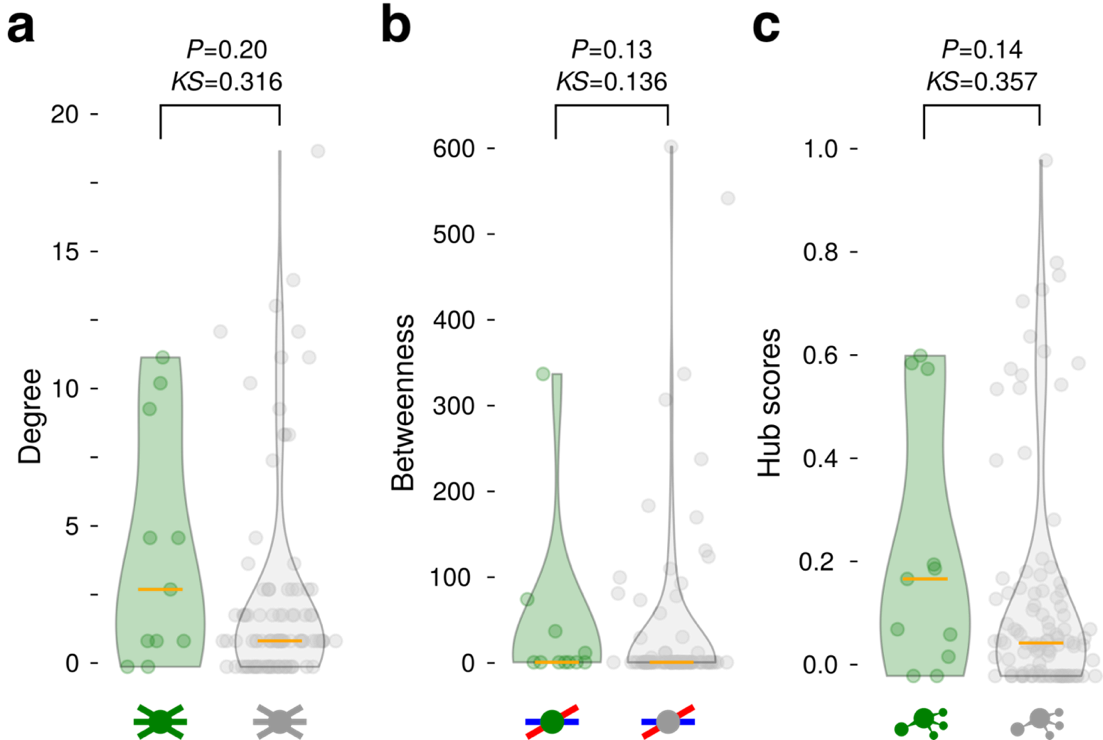

**Fig. 5. Simple centrality measures show no difference between cores and other neurons**. **a**, The degree – number of connections per neuron – of cores was not significantly higher than other neurons. **b**, The betweenness – the number of shortest paths between any two nodes passing by a considered node – of cores was not significantly different from other neurons. **c**, The hub score – the weighted number of outward connections of a node that points to central nodes – of cores was not significantly higher than other neurons.

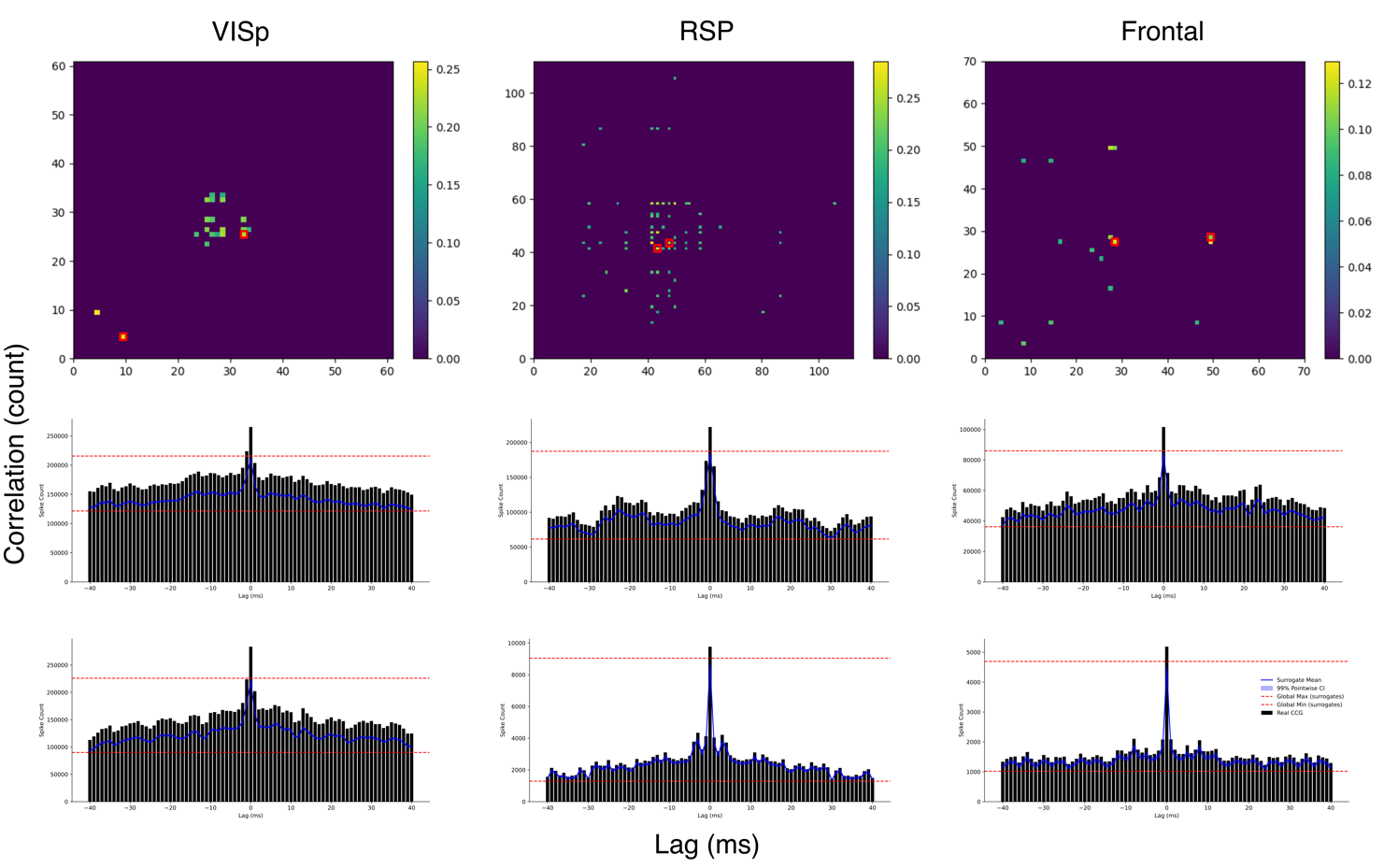

**Fig. 6. Cross-correlograms of Neuropixels units.** Cross-correlation was computed over correlated units from the Neuropixels datasets for the areas of interest (top, population correlograms, red squares are the units correlograms shown below). Correlograms were computed as in Fujisawa et al. 2008. Briefly, spike trains from pairs of units were convolved using bins of 2ms over a window of 40ms (black bars). Then, each spike train was independently jittered within ±5 ms to generate 1000 surrogate datasets. The surrogate mean (blue curve), 99% pointwise (blue shaded area) and global confidence bands (red lines) are derived from the surrogate distributions. This analysis informed our choice of the time window (1-5ms) to estimate monosynaptic connections in our functional connectivity analysis (see Methods).

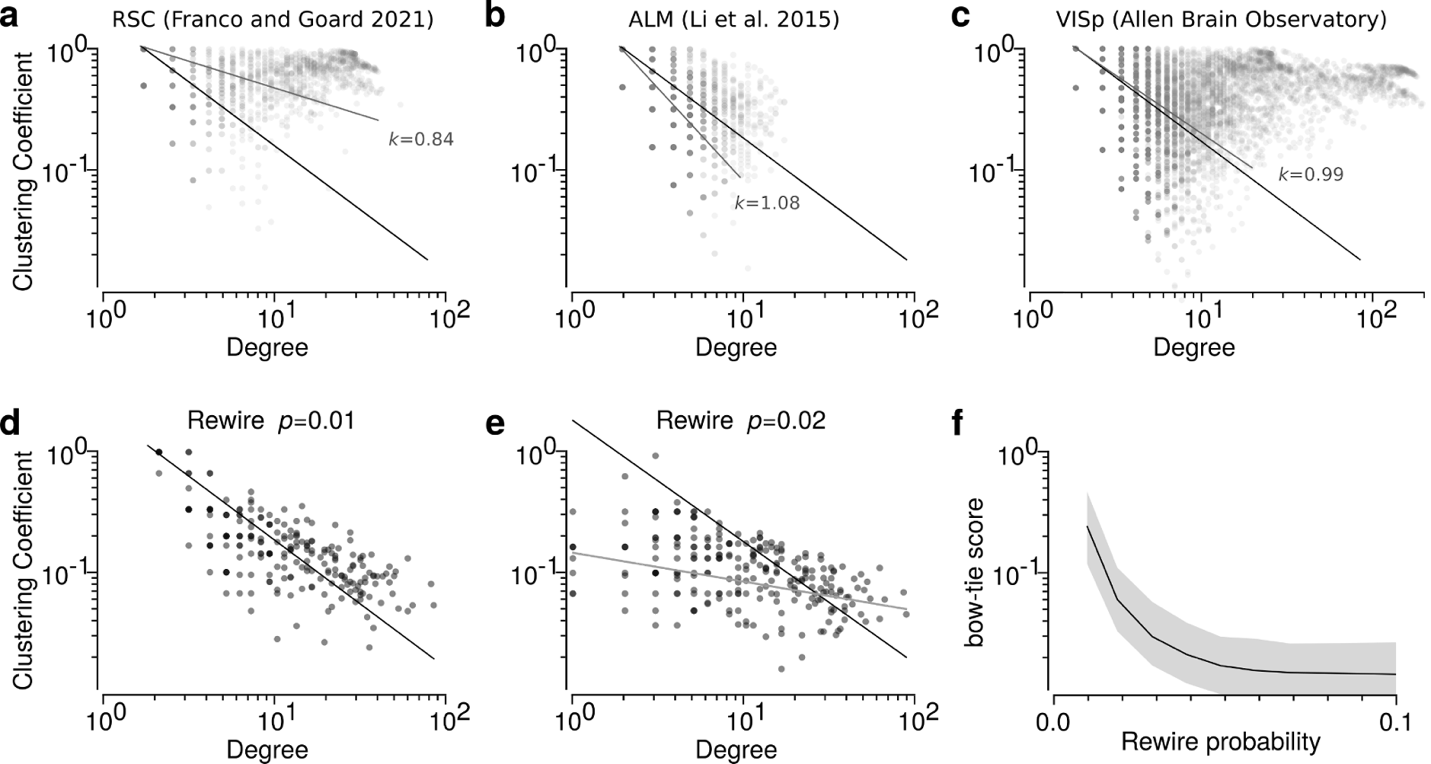

**Fig. 7. Hierarchical modularity across cortices and rewiring of the MICrONS dataset**. **a**-**c**, Example hierarchical modularities for other cortices (Retro-Splenial Cortex, from Franco and Goard 2021, and Antero-Lateral Motor cortex, from Li et al. 2015), computed from functional activity using the method described by Sadovsky and MacLean 2013. Each cortical region had a different slope (in black the fitting curve for the MICrONS data). This modularity, estimated using functional connectivity, may tend to overestimate large degrees / large clustering coefficient, as evident comparing the same area (VISual primary) functional hierarchical modularity of Fig. 3f and h. **d** and **e**. Example hierarchical modularity resulting from rewiring the MICrONS graph. **d**, rewiring the graph edges with probability *p*=0.01 results in a hierarchically modular relationship as that observed in the data. In **e**, already at *p*=0.02 the relationship deteriorates (fitting curve in black is kept in both panels). **f**, The bow-tie score (ratio of modules with clearly identifiable submodules converging-diverging from a central module over the total number of modules) rapidly deteriorates for increasing rewiring probabilities (SEM reported as grey shaded area).

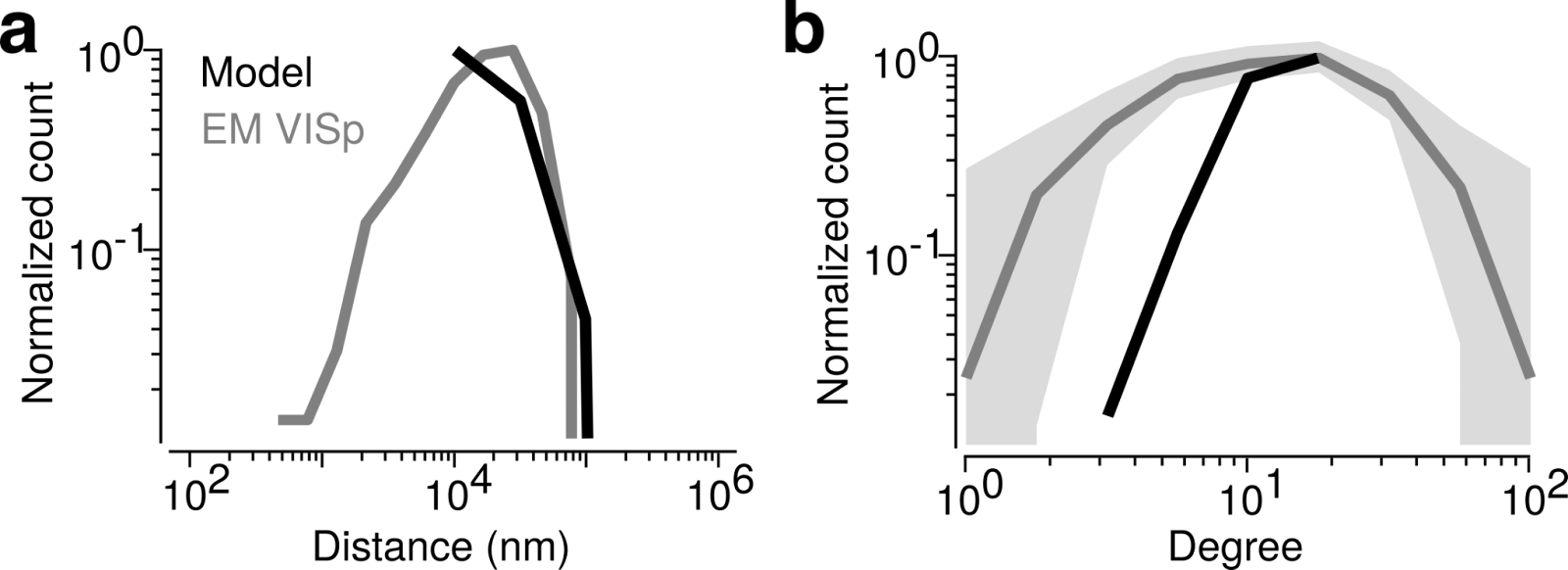

**Fig. 8. Comparison of the MICrONS and model basic network properties**. **a**, Distribution of distances between connected neurons in the MICrONS dataset (grey). Notice that its shape follows the ones found in Kätzel et al. 2011, Hage et al. 2022, Seeman et al. 2009, and on a larger scale by Gamanut et al. 2018. We fitted our model (black) to the data, considering the space available given the inter-cell minimal distance assumed as model spatial and density resolution (1 cells / 100 µm^2^, for a total of 10000 excitatory cells/mm^2^, 2500 inhibitory cells are distributed accordingly to span the same space). **b**, Normalised count of degrees in the MICrONS dataset (grey, with shaded SEM). In the model (black), the max degree is 75, which we imposed in all models. For small distance-dependent connectivity ranges (from 22 to 42 µm) the chosen inter-cell minimal distance resulted in a lower number of connections in the model.

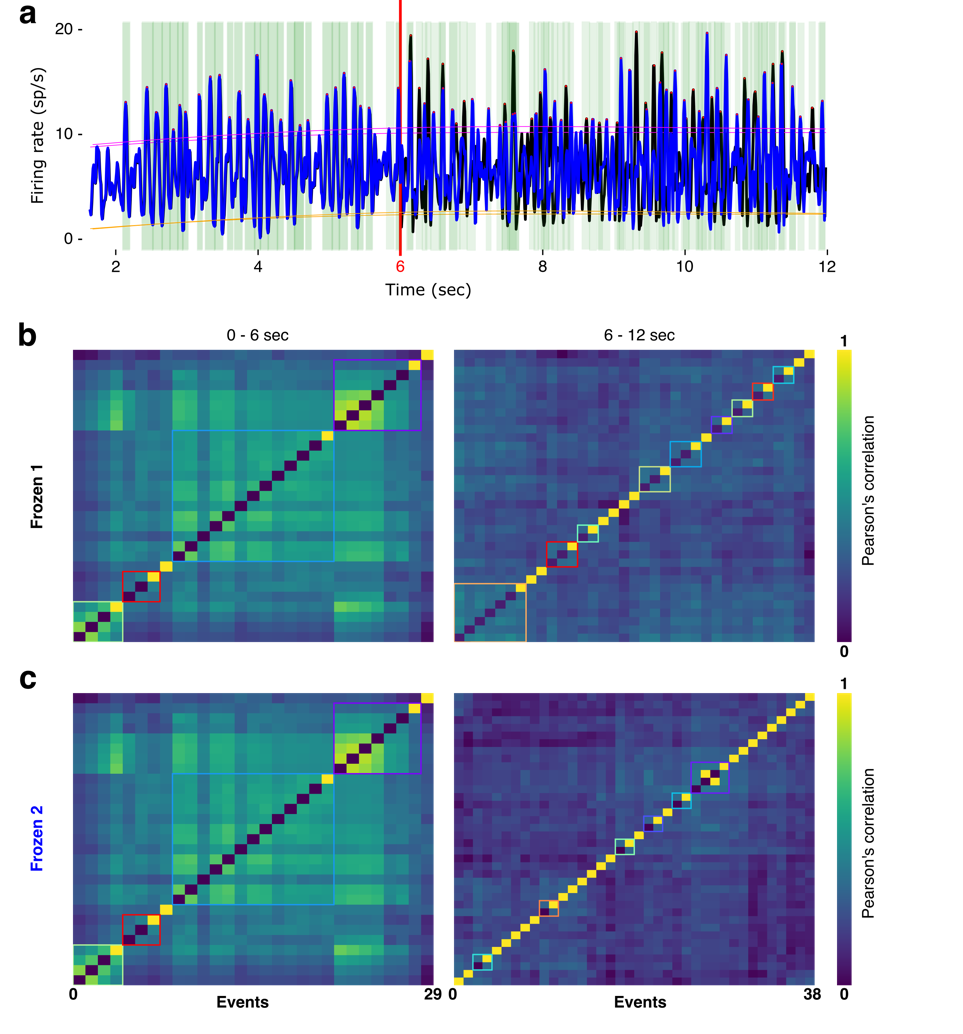

**Fig. 9. Specific clusters for different stimuli.** We tested the ability of our mechanistic model to support pattern completion to show that it can recapitulate phenomena in V1 that have been specifically cited as evidence that it is an attractor network. In many previous models, fixed-point attractors (pattern completion subnetworks) are achieved either through learning or artificially imposed. In our model no learning nor artificial subnetwork was imposed, yet reproducible firing patterns are observed. We used a “frozen noise” input strategy (Marre et al. 2009) to see whether it leads to same pattern completions. Three spike train matrices (A, B, C) were generated using Poisson populations with different seeds (all having on average 5 sp/s). We then created two spike trains matrices (“Frozen 1” and “Frozen 2”) in which the first half was identical for both (A), and the second half varied (either B or C). Then we ran the same model as in Fig. 5 using the two (frozen) spike train matrices to test whether it reproduced the same events during the identical first half, and different events in the second half. **a**, Firing rates from the network stimulated with the two spike train matrices (“Frozen 1”, black, and “Frozen 2”, blue). The first portion of the firing (up to 6sec, red vertical line) is the same for both runs. The second portion of the firing rate was different (green vertical shades: synchronous events). **b-c**, The clustering of population events shows the complete reproducibility of activity for the initial portion (first panel in **a** and **b**). Different clusters emerged in the second portion of the stimulus (second panel in **a** and **b**) (colored squares: clusters of vectors sharing neurons beyond a surrogate-based threshold).

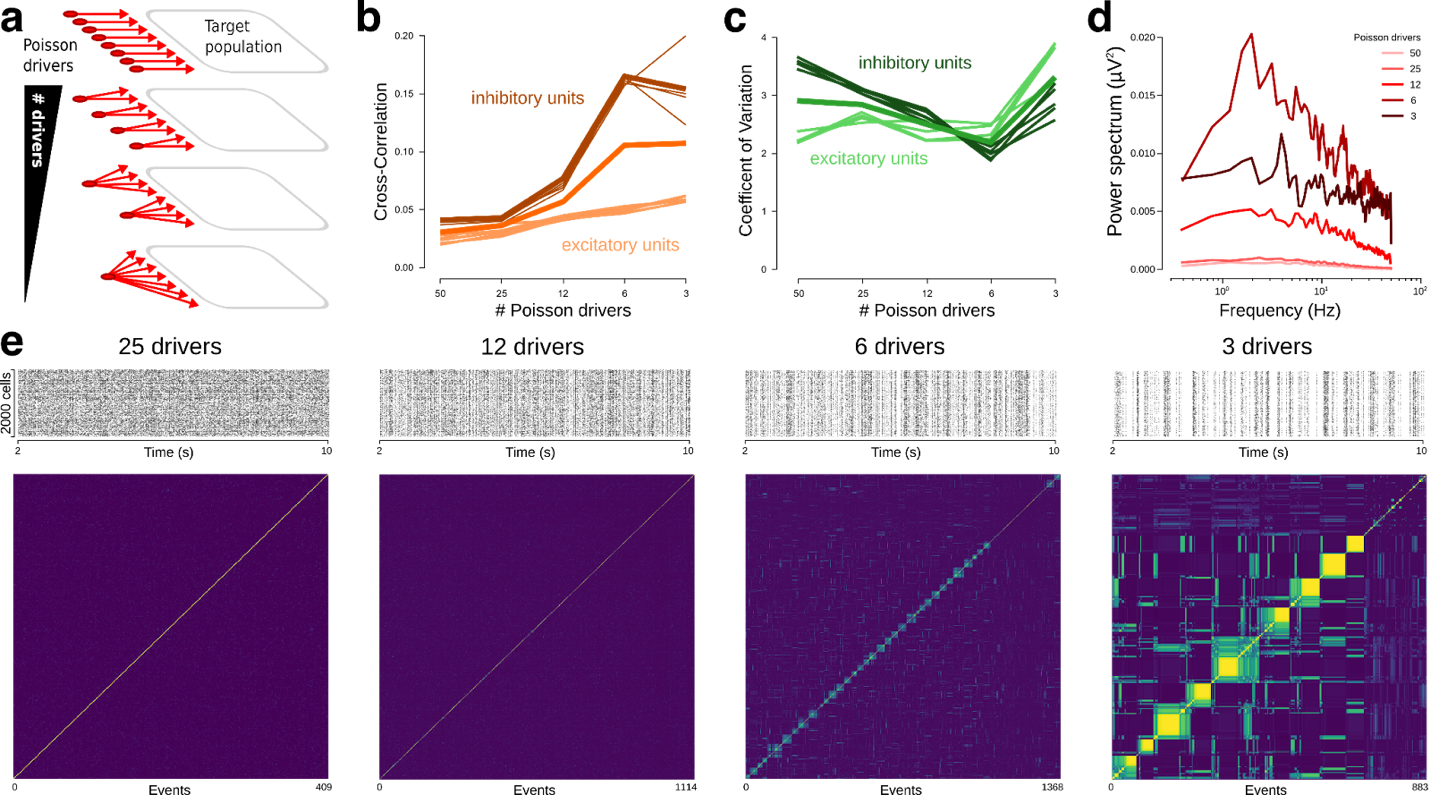

**Fig. 10. Increasing input correlations rescues population event reproducibility but introduces network-wide oscillations**. **a**, For the same target population (right), we systematically reduced the number of Poisson input drivers (red disks and arrows), to explore how the increase in input correlations affected the correlations between population events. **b**, Reducing the number of Poisson input drivers (n=[50, 25, 12, 6, 3]) increased the population cross-correlation of spikes toward values characteristic of oscillating regimes (from CC=0.035 at n=50, to CC=0.11 at n=3). **c**, The number of Poisson drivers did not alter significantly the population coefficient of variation. **d**, Reducing the number of Poisson input drivers increased the power for slow oscillations in the Fourier spectrums of the firing rates. **e**, Four 10 sec spike rasters for 2000 example cells, with their correlation matrices, show the changes in firing regimes responsible for the increase in population events correlation. The first correlation matrix on the left shows the absence of reproducibility. In the right correlation matrix, the off-diagonal correlations show the presence of correlations between population events, resulting in higher values of reproducibility.
